# Seasonal influenza circulation patterns and projections for September 2019 to September 2020

**DOI:** 10.1101/780627

**Authors:** Trevor Bedford, John Huddleston, Barney Potter, Richard A. Neher

## Abstract

This report details current seasonal influenza circulation patterns as of August 2019 and makes projections up to September 2020 to coincide with selection of the 2020 Southern Hemisphere vaccine strain. This is not meant as a comprehensive report, but is instead intended as particular observations that we’ve made that may be of relevance. Please also note that observed patterns reflect the GISAID database and may not be entirely representative of underlying dynamics. All analyses are based on the nextflu/nextstrain pipeline [1,2] with continual updates posted to nextstrain.org/flu. **A/H3N2:** A/H3N2 viruses continue to show substantial diversity in HA sequences with a deep split between 3c3.A and 3c2.A1b viruses. The most notable recent developments are the rapid rise of clade A1b/137F – a subclade of A1b/135K – in China and Bangladesh and clade A1b/197R – a subclade of A1b/131K – which dominates the ongoing season in Australia. Our models predict that A1b/137F and A1b/197R will be the dominant clades next year with A1b/197R accounting for most circulation. There is, however, large uncertainty in the true extent of A1b/137F circulation. **A/H1N1pdm:** The S183P substitution has risen to near fixation. The most successful subclade carrying this mutation is 183P-5 which has essentially replaced competing variants. A variant with substitutions 129D/185I is at 60% prevalence globally, while a second variant with substitution 130N is at 50% in North America and ~10% elsewhere. Substitutions at site 156 to D or K have arisen sporadically and result in loss of recognition by antisera raised against viruses with asparagine at position 156. Despite the large antigenic effect, viruses with mutations at site 156 don’t seem to spread. Beyond variants at site 156, little to no antigenic evolution is evident in assays with ferret antisera. **B/Vic:** Antigenically drifted deletion variants at HA1 sites 162, 163 and 164 are now dominating global circulation and have all but taken over. The double deletion variant V1A.1 had previously been circulating at high frequency in the Americas. However, over the course of 2009, the triple deletion variant V1A.3 has increased in frequency globally and is now dominating in all geographic regions. Importantly, V1A.1 and V1A.3 variants appear antigenically distinct by HI assays with 4-8 fold reductions in log2 titer in both directions. **B/Yam**: B/Yam has not circulated in large numbers since the Northern Hemisphere season 2017/2018 and displays relatively little amino acid variation in HA or antigenic diversity. Amino acid variants at sites 229 and 232 have begun to circulate and population is now split between 229D/232D, 229N/232D and 229D/232N variants. These variants show little sign of antigenic difference in HI assays.

## Methods and Notes

### Sequence data and subsampling

We base our analysis on sequence data available in GISAID as of Sep 16, 2019 [3]. The availability of sequences varies greatly across time and geography and we try to minimize geographical and temporal bias by subsampling the data or analyzing different geographical regions separately when appropriate. While this subsampling reduced geographical biases, it doesn’t remove this bias entirely. The most recent months are particularly prone to biases due to variable data deposition schedules across geographic regions.

### Phylogenetic analysis

The database contains too many sequences to perform a comprehensive phylogenetic analysis of all available data. We hence subsample the data to 90 sequences per month using the following criteria:

- for each month and each of ten geographic regions, we select 9 viruses (or all available viruses if fewer than 9 are available);
- when the total number of viruses selected by the above criterion in a given month is below 90, we fill the remainder evenly with viruses from geographic regions with more available strains;
- within each month and geographic region, viruses with antigenic data are prioritized.

Parallel evolution, that is repeated occurrence of identical substitutions in different clades of the tree, has become common in A/H3N2 and A/H1N1pdm. Such parallel evolution violates fundamental assumptions of common phylogeny software and can erroneously group distinct clades together if they share too many parallel changes. To avoid such artifacts, we mask sites with rampant parallelism prior to phylogeny inference. Clades are assigned using a collection of “signature” mutations available at the GitHub repository github.com/nextstrain/seasonal-flu for each lineage via the following links:

- config/clades_h3n2_ha.tsv
- config/clades_h1n1pdm_ha.tsv
- config/clades_vic_ha.tsv
- config/clades_yam_ha.tsv

### Mutation frequency calculations

In contrast to the phylogenetic analysis, mutation frequencies are based on all available data and calculated separately for each geographic region. To obtain estimates of global mutation frequencies, we average geographic regions weighted by their approximate contribution to the global human population. Specifically, frequencies are calculated as follows:

- amino acid sequences of all isolates within a geographic regions are aligned to a reference sequence;
- for each variable alignment column, we infer a frequency trajectory of the different amino acids at this position using a Brownian motion prior [1];
- in each region, the seasonal pattern of sequence data availability is used as a proxy for seasonal prevalence;
- regional frequencies are then averaged both by the seasonal pattern and their population fraction.

The graphs in the report show frequency trajectories for North- and South America, China, Japan/Korea, Europe, and Oceania and omit Africa, South Asia, West Asia, and Southeast Asia to avoid overloading the graphs although these regions still contribute to globally weighted estimates.

### Antigenic analysis

We summarize HI and FRA measurements provided by the WHO CCs in London, Melbourne, Atlanta and Tokyo using our *substitution model* [4] which models log-titers as a sum of effects associated with amino acid differences between the sequences of the test and reference virus. In addition, the model allows for a serum (column) and a virus (row) effect. This model allows to infer titers for virus/serum pairs that have not been antigenically characterized and isolates effects consistently observed across many measurements from the noise inherent in individual measurements.

### Persistent analyses

For a better historical record, the analyses available on Sep 23, 2019 have been saved as:

- nextstrain.org/flu/seasonal/h3n2/ha/2y/2019-09
- nextstrain.org/flu/seasonal/h1n1pdm/ha/2y/2019-09
- nextstrain.org/flu/seasonal/vic/ha/2y/2019-09
- nextstrain.org/flu/seasonal/yam/ha/2y/2019-09

## A/H3N2

A/H3N2 viruses continue to show substantial diversity in HA sequences with a deep split between 3c3.A and 3c2.A1b viruses. Clades A2/re, A3 and A4 are now rare. 3c3.A had been observed at low levels throughout the last four years in the Western Hemisphere and rose to high frequencies in the 2018/2019 season before falling again in frequency recently. It remains geographically restricted to the Americas and isolated European countries. The substantial antigenic distance between 3c3.A and A1b poses a threat of vaccine mismatch. The most notable recent developments are the rapid rise of clade A1b/137F – a subclade of A1b/135K – in China and Bangladesh and clade A1b/197R – a subclade of A1b/131K – which dominates the ongoing season in Australia. Our models predict that A1b/137F and A1b/197R will be the dominant clades next year with A1b/197R accounting for most circulation. There is, however, large uncertainty in the true extent of A1b/137F circulation.

We base our primary analysis on a set of viruses collected between Sep 2017 and Aug 2019, comprising upwards of 300 viruses per month in the Northern hemisphere winter but fewer counts otherwise (Fig. 1).

**Figure 1.**
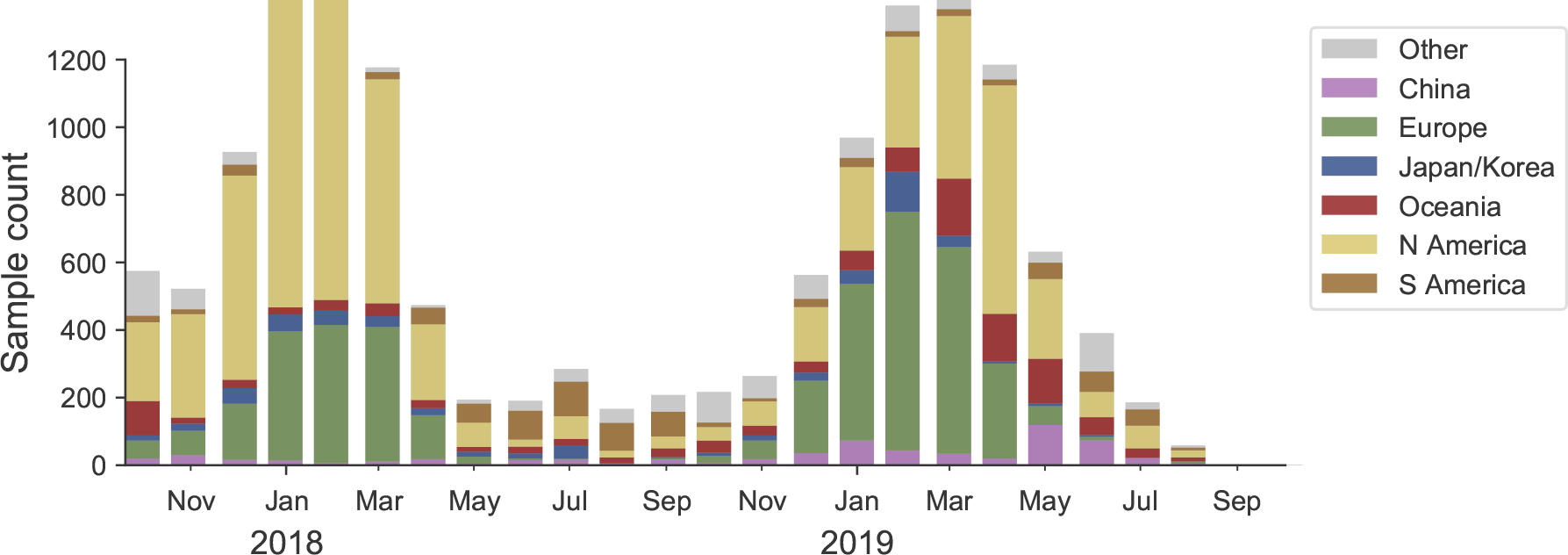
Sample counts through time and across regions. This is a stacked bar plot with the visible height of a color bar corresponding to the sample count from the respective region.

### Current circulation patterns

Over the past three years, 3c2.A viruses within A/H3N2 have differentiated into multiple major cocirculating clades with variable geographic distributions (Fig. 2). The past year was dominated by clade A1b. The reassortant clade A2/re that was common in the NH 2017/2018 is now rare. Subclades A1b/131K and A1b/135K comprised the majority of A1b viruses. Over the last 18 months, 3c3.A viruses have increased markedly in the US and Europe and accounted for 60% of isolates in North America and 10-20% in Europe during the last NH winter, see Fig. 4.

**Figure 2.**
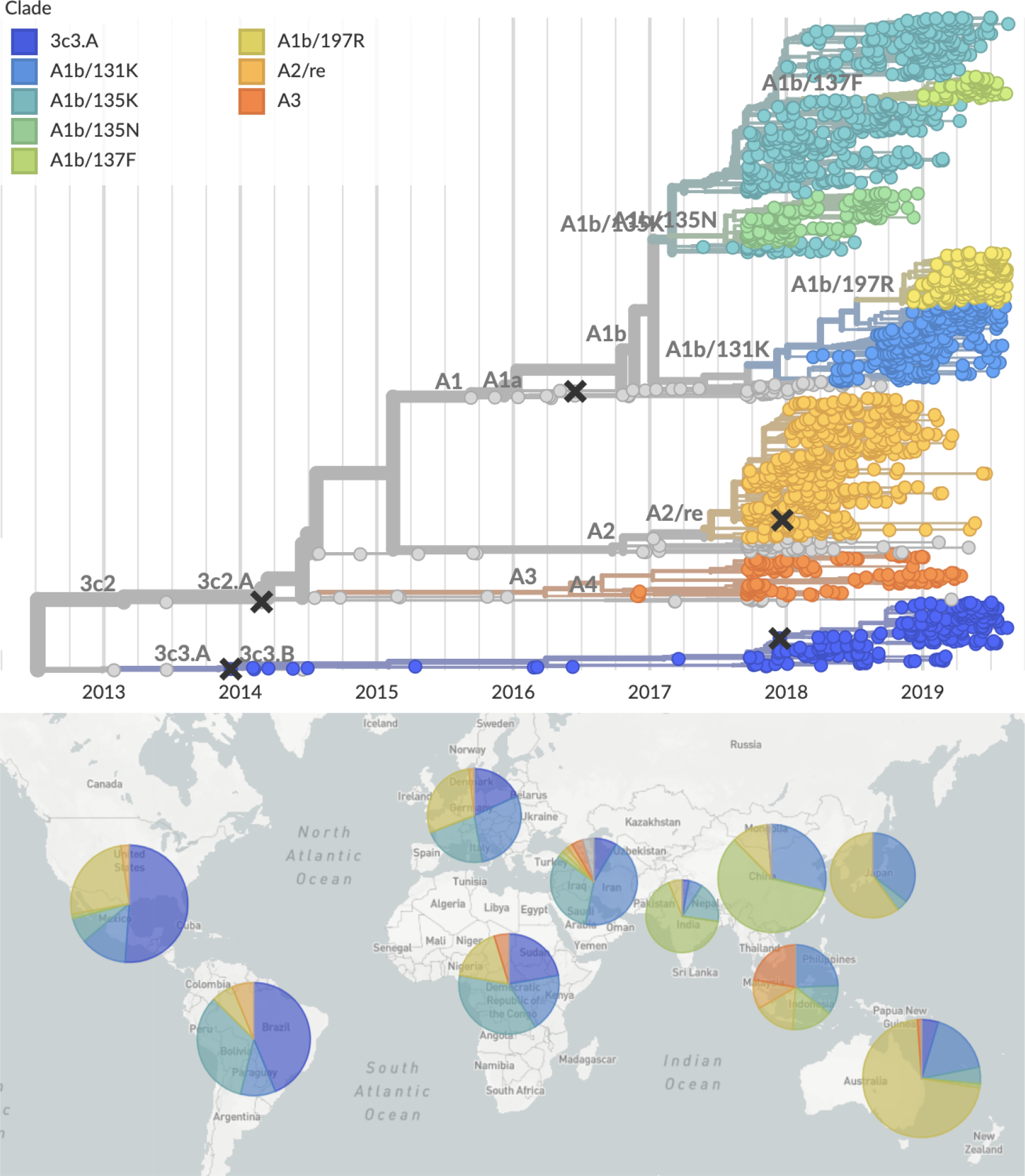
H3N2 phylogeny colored by clade and the corresponding geographic breakdown. The shown geographic distribution is for viruses from 2019 only.

Recently, new subclades within A1b have emerged and risen to appreciable global frequency. These are A1b/197R that has appeared on top of the A1b/131K background and A1b/137F that has appeared on top of the A1b/135K background. A1b/197R viruses have predominated the 2019 SH Oceania season and are now commonly observed in the NH as well. A1b/137F viruses have largely been localized to China and Bangladesh, but are prevalent within these regions. Outside of Bangladesh, very few sequences from South Asia (India in particular) have been deposited in GISAID. There is therefore substantial uncertainty in the estimates of the total circulation of A1b/137F viruses.

Major clades and their associated substitutions include:

- Clade A1b/131K: 142G, T131K
- Clade A1b/135K: 142G, T135K
- Clade A1b/135N: T135N
- Clade A1b/137F: T135K, T128A, S137F, A138S, F193S (subclade within A1b/135K)
- Clade A1b/197R: T131K, Q197R, S219F (subclade within A1b/131K)
- Clade A2/re: T131K, R142K, A212T (reassortant clade)
- Clade 3c3.A: S91N, N144K, F159S, F193S

The frequency trajectories of the these major clades in different geographic regions are shown in Figure 4. In the first half of 2018, clade A2/re and A1b/135K were dominant. Subsequently subclade A1b/131K rose in frequency towards the end of 2018 and now accounts for 50% of global circulation with an increasing trend. Clade A1b/131K and subclade A1b/197R have dominated the anomalously early season in Australia. Clade A1b/135K has had a wide geographic distribution for the last 2 years and stayed at a steady frequency of about 30% globally, albeit with changing regional prevalence. The subclade A1b/137F with additional substitutions A138F and F193S has recently become common in China and Bangladesh and is expanding rapidly. The clade A2/re has not been observed in recent months with the exception of sporadic isolation in South America. Clade 3c3.A continues to be common in the Americas, while having been partly replaced by A1b/131K in North America over the past couple of months. These dynamics of clade frequencies are reflected in the local branching index and other measures of recent clade growth discussed later in the report (see Figure 3).

**Figure 3.**
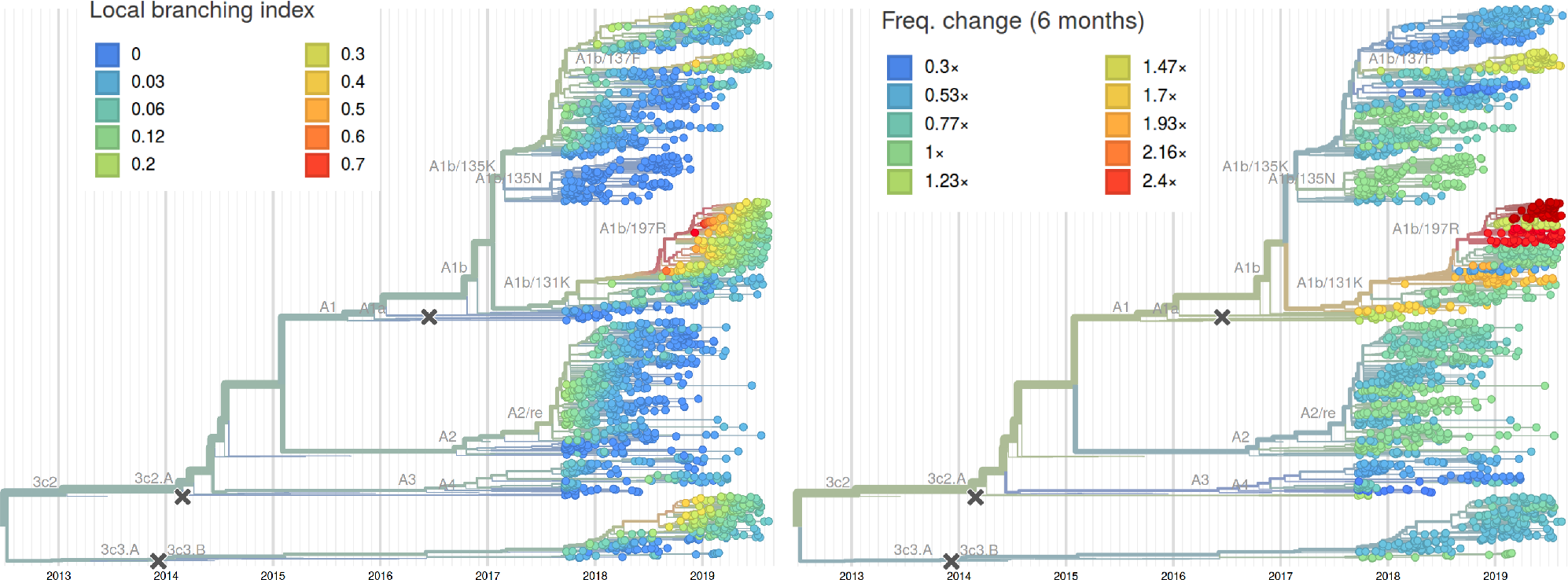
H3N2 phylogenies colored by growth. Left: Colored by LBI [5], a measure of recent clade expansion that is predictive of future success. Right: Colored by recent clade fold-change in frequency. Clade A1b/197R scores highest on both measures. The newly emerging clade A1b/137F scores second highest for recent clade growth, while 3c3.a and A1b/137F are tied for LBI. The LBI averages clade growth over longer time and factors in frequency information, while the growth measure on the right is more sensitive to recent changes and rapid growth of small clades.

**Figure 4.**
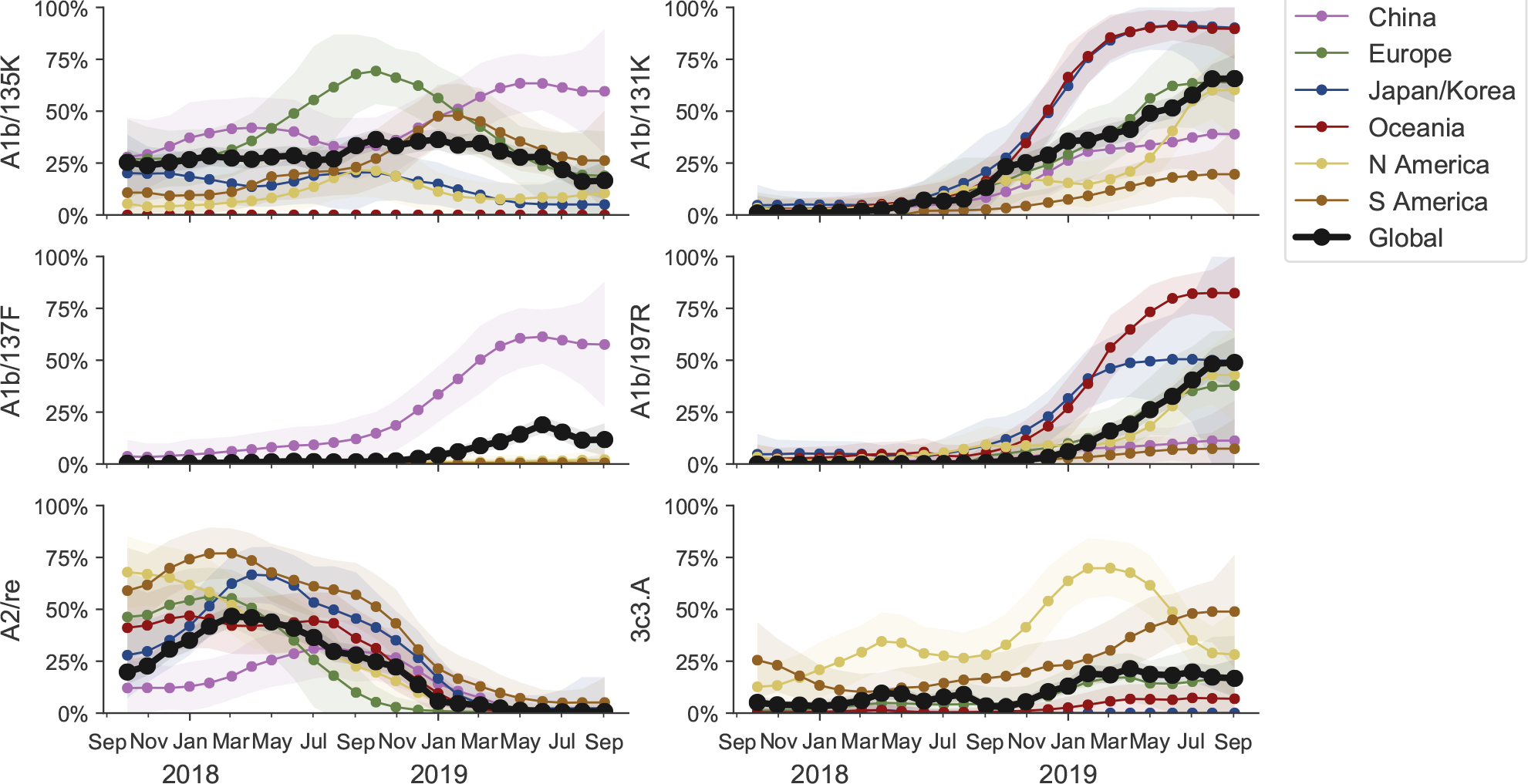
Frequency trajectories of H3N2 clades partitioned by clade then by region. We estimate frequencies of different clades based on sample counts and collection dates of strains included in the phylogeny. We use a Brownian motion process prior to smooth frequencies from month-to-month. Transparent bands show an estimate of the 68% confidence interval based on sample counts.

Frequencies of specific substitutions are shown in Figure 5. We would like to draw attention to the following patterns of convergent evolution and recent emergence:

- **HA1:T128A** arose in a subclade of **A1b/135K** and clade **3c3.A**, but not in A1b/131K.
- **HA1:T131K** was common in the 2017/2018 season as part of **A2/re** and rose again in frequency as part of clade **A1b/131K**
- **HA1:T135K** has arisen multiple times in the recent past, but its current circulation is restricted to clade **A1b/135K**.
- **HA1:137F** recently emerged along with 138S and 193S and rose rapidly in China to levels above 60%. Recent viruses sampled in Bangladesh also fall into this clade, but it has only been observed sporadically elsewhere. **3c3.A** also has A138S.
- **HA1:142G** arose in clades **A1b/135K**, **A1b/131K**, and **3c3.A** and has replaced all other variants at this position.
- **HA1:197R** arose within clade A1b/131K and is dominating the season in Australia. This substitution sits on a 219F background.

**Figure 5.**
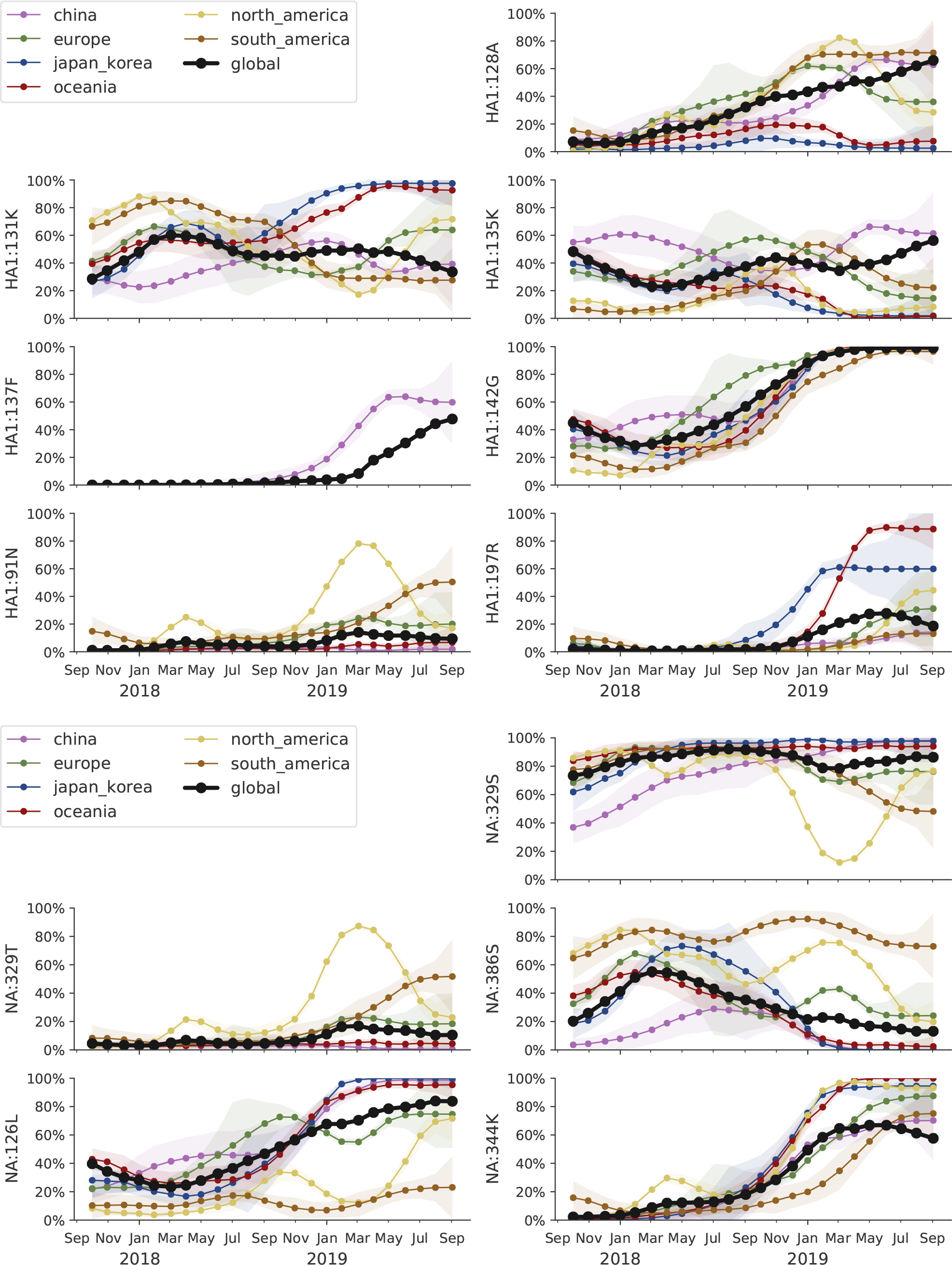
Frequency trajectories of substitutions in H3N2 HA and NA. We estimate frequencies of different amino acid variants based on sample counts and collection dates. These estimates are based on all available data. We use a Brownian motion process prior to smooth frequencies from month-to-month. Transparent bands show an estimate of the 68% confidence interval based on sample counts.

### Antigenic properties

We integrated HI and FRA/VN data from the WHO Collaborating Centers in London, Tokyo, Melbourne and Atlanta with molecular evolution of the HA segment. Patterns of recent antigenic evolution are most clearly seen in FRA titers using serum raised against A/NorthCarolina/4/2016 (see Fig. 6). Our titer model suggests a small drop (2-fold) in titers for clade A1b/197R and a moderate drop (2- to 4-fold) for clade A1b/137F. The latter inference, however, is based on a small number of measurements. Overall, little antigenic evolution is detectable by ferret antisera within the 3c2.A clade despite frequent and recurring changes in at epitope sites. Clade 3c3.a, however, is clearly antigenically distinct.

**Figure 6.**
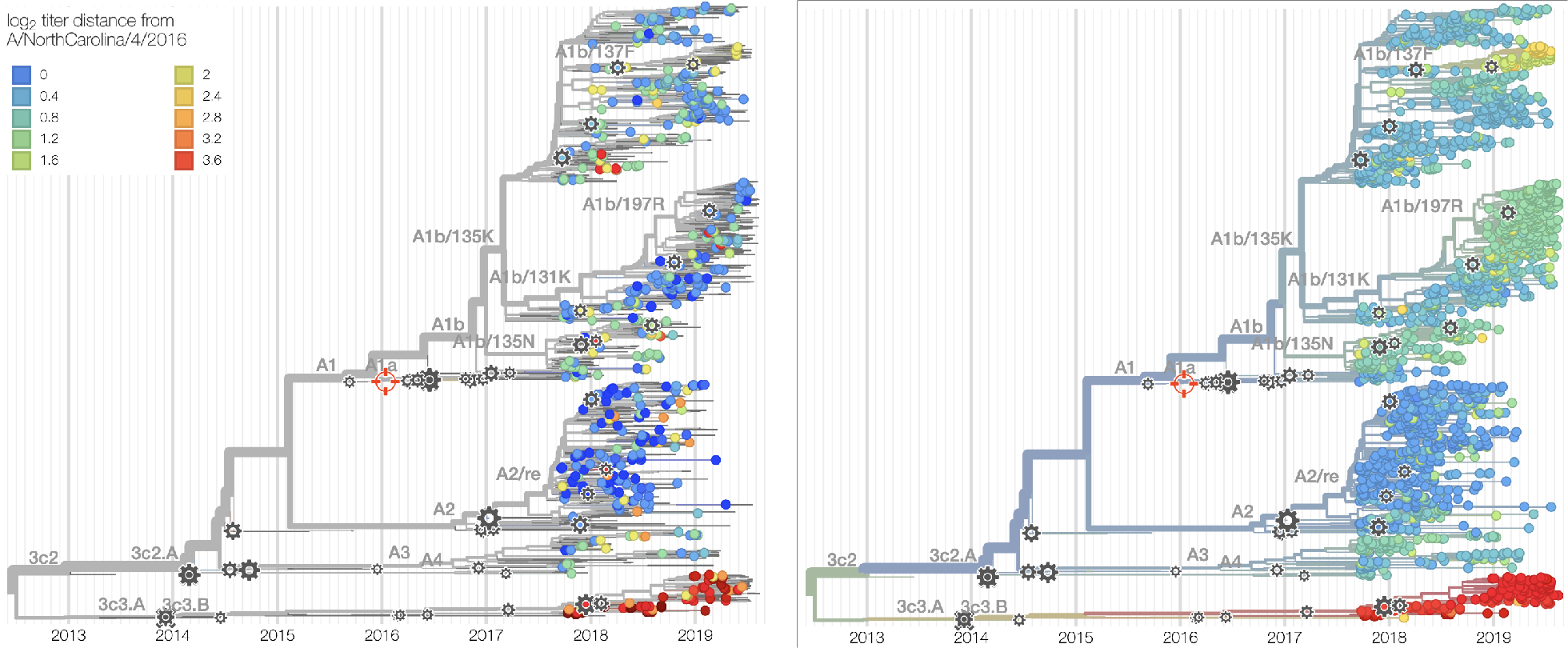
H3N2 phylogeny colored by antigenic distance as measured by FRA to A/NorthCarolina/4/2016. The left panels show the individual measurements, the right panel the model inference. Blue corresponds to good coverage, yellow to a 4-fold drop, red to a 16-fold drop (or higher).

### Fitness model

Here, we construct a fitness model to estimate fitness of circulating virus strains and project strain and clade frequencies. We take an approach of measuring an assortment of metrics that may correlate with fitness and then using historical data to weight fitness components in a combined model. We assess the degree to which we can predict changes in strain frequency in one year lookaheads, e.g. how well do projections in Feb 2017 predict strain frequencies in Feb 2018?

Generally, we combine a metrics for antigenic novelty, intrinsic fitness and recent clade growth. More specifically, we include cross-immunity mediated by substitutions at epitope sites, cross-immunity mediated by HI titer differences, substitutions at non-epitope sites as an inverse proxy for intrinsic fitness and recent clade growth as measured by local branching index (LBI). However, in recent years we only observe consistent performance from local branching index with a small improvement from the inclusion of non-epitope sites (Fig. 7). Measurements of antigenic novelty either by epitope mutations or by HI titers have not consistently improved model performance since 2009. Thus, we believe the most robust model is one that focuses on LBI supplemented by non-epitope sites. Here, we present results from an LBI-only model as well as a model that combines LBI and non-epitope mutations.

**Figure 7.**
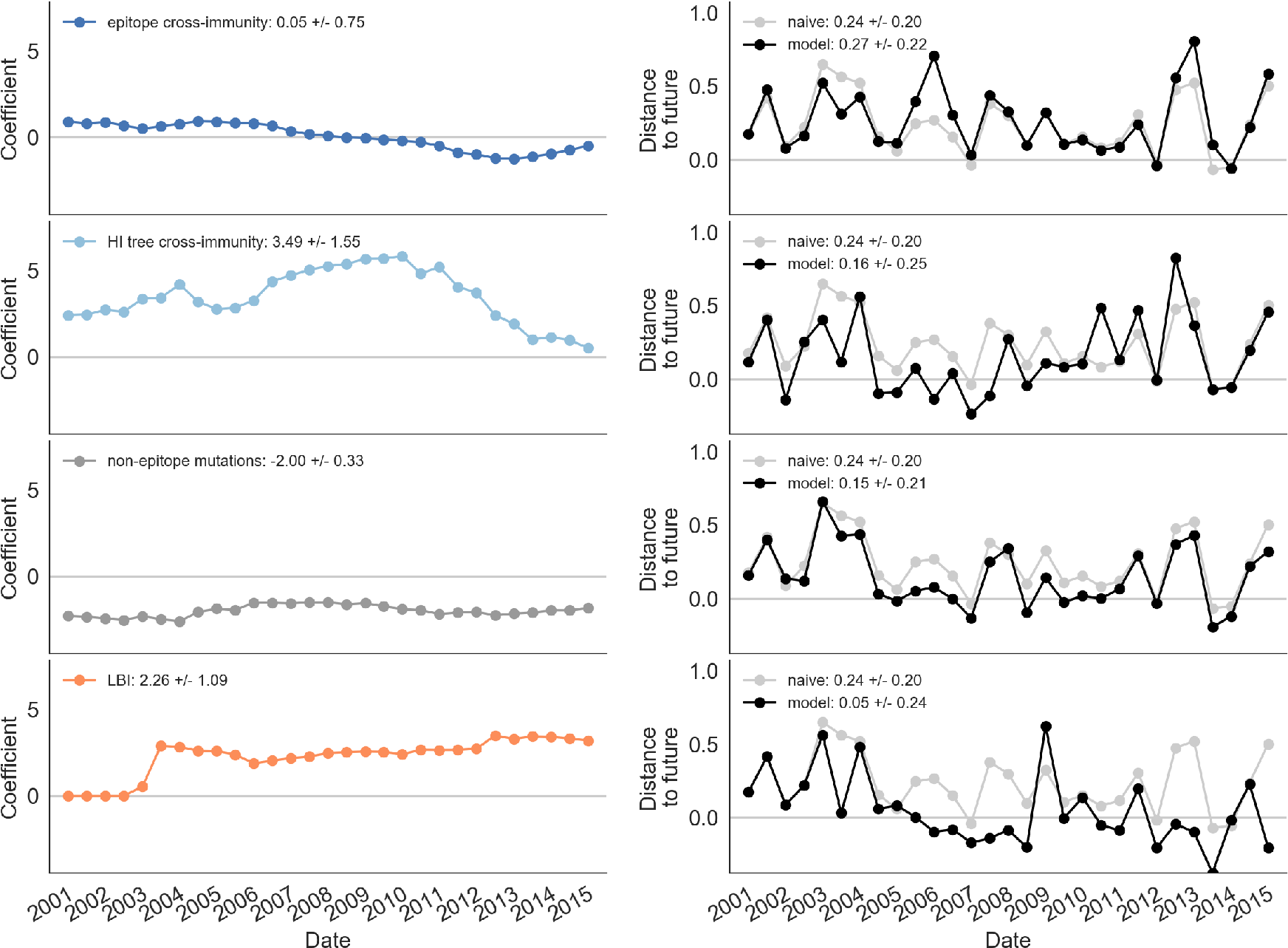
Model performance in previous seasons. Left-hand panels show predictor coefficients; a good predictor will have a consistent non-zero coefficient through time. Right-hand panels show model performance as Hamming distance between predicted strain distribution and retrospectively observed strain distribution; a good model will have black lines (showing model predictions) consistently below gray lines (showing predictions of a ‘naive’ model).

As illustrated previously (Fig. 3), clade A1b/197R viruses have been increasing in frequency and have the highest values of local branching index. This results in clade A1b/197R and clade A1b/131K having higher fitness than A1b/135K, A1b/137F and 3c3.A viruses. We use these estimated strain fitnesses to project frequencies of individual strains and from these projections forecast resulting clade frequencies (Fig. 8). This shows that both the LBI-only and the LBI with non-epitope sites predictions favor A1b/197R viruses to grow in frequency from ~32% global frequency to ~80% global frequency over the next 12 months. This projection has A1b/131K viruses expanding but due completely to the success of A1b/197R. A1b/137F are predicted to persist into the future but at lower frequency than A1b/197R viruses.

**Figure 8.**
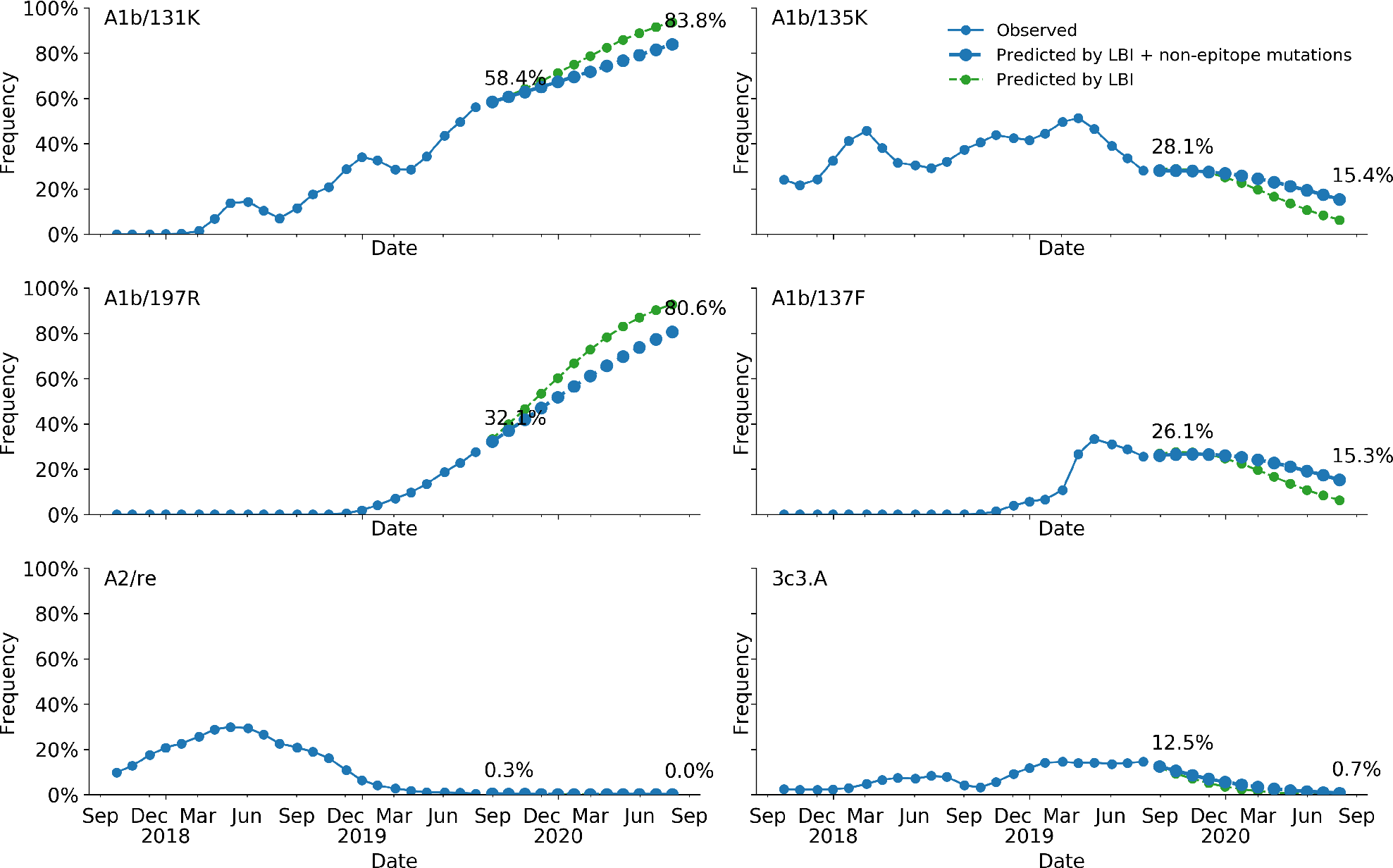
Frequency projection based on fitness model. Having assigned each virus strain with an estimated fitness, it’s possible to project clade dynamics. Projections are shown for models using only single predictors as well as for the combined composite model.

It is important to point out that it is likely that clade A1b/197R is better sampled than A1b/137F. Since the LBI is effectively a composite measure of frequency and recent expansion, sampling bias might have led to a too high fitness estimate of A1b/197R and a too low estimate for A1b/137F. Both of these clades will likely persist at high frequencies.

## A/H1N1pdm

The S183P substitution has risen to near fixation. The most successful subclade carrying this mutation is 183P-5 which has essentially replaced competing variants. This subclade P5, however, is in itself deeply split with a geographically heterogeneous distribution. A variant with substitutions 129D/185I is at 60% prevalence globally, while a second variant with substitution 130N is at 50% in North America and ~10% elsewhere. Substitutions at site 156 to D or K have arisen sporadically and result in loss of recognition by antisera raised against viruses with asparagine at position 156. Despite the large antigenic effect, viruses with mutations at site 156 don’t seem to spread. Beyond variants at site 156, little to no antigenic evolution is evident in assays with ferret antisera.

We base our primary analysis on a set of viruses collected between Jul 2017 and Aug 2019, comprising upwards of 100 viruses per month in almost all months (Fig. 9). We use all available data when estimating frequencies of mutations and weight samples appropriately by regional population size and relative sampling intensity to arrive at a putatively unbiased global frequency estimate. Phylogenetic analyses are based on a representative sample of about 2000 viruses.

**Figure 9.**
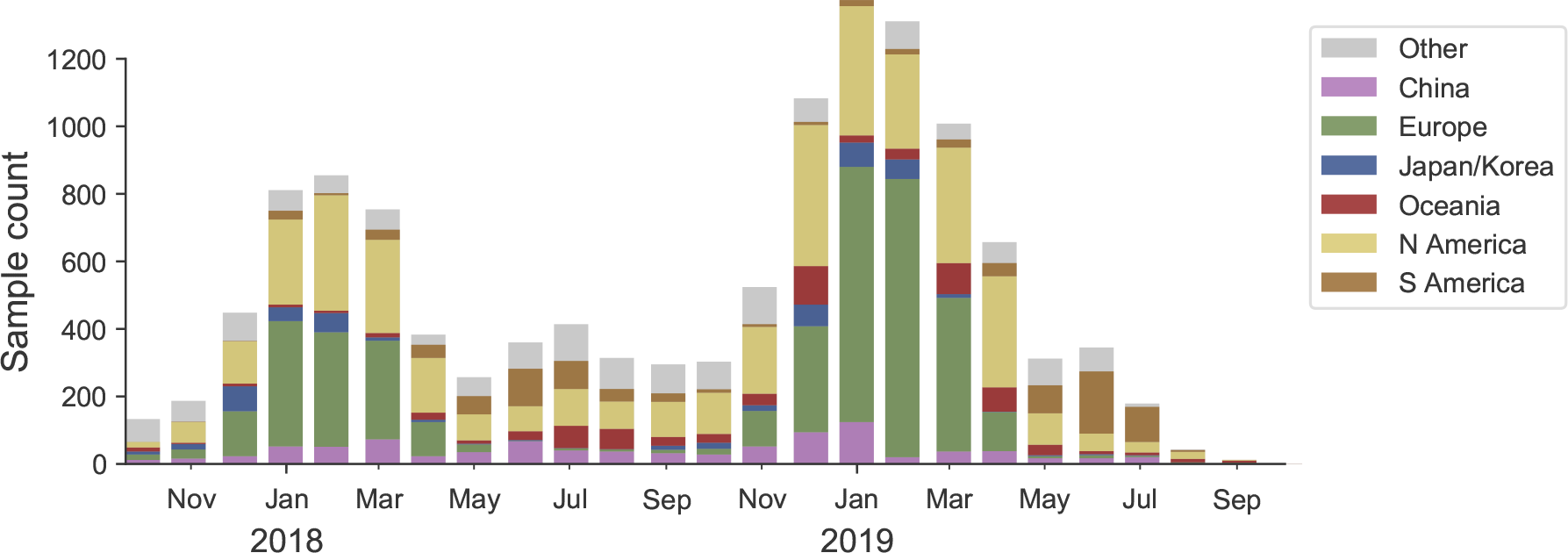
Sample counts through time and across regions. This is a stacked bar plot with the visible height of a color bar corresponding to the sample count from the respective region.

Over the course of the last 2 years, the substitution S183P has arisen multiple times and viruses carrying this substitution have almost completely taken over (Fig. 10). Of the main clades carrying the 183P substitution (labeled P1-P7), clade P5 accounts for 70-100% of recent circulation in different geographic regions (Fig. 12).

**Figure 10.**
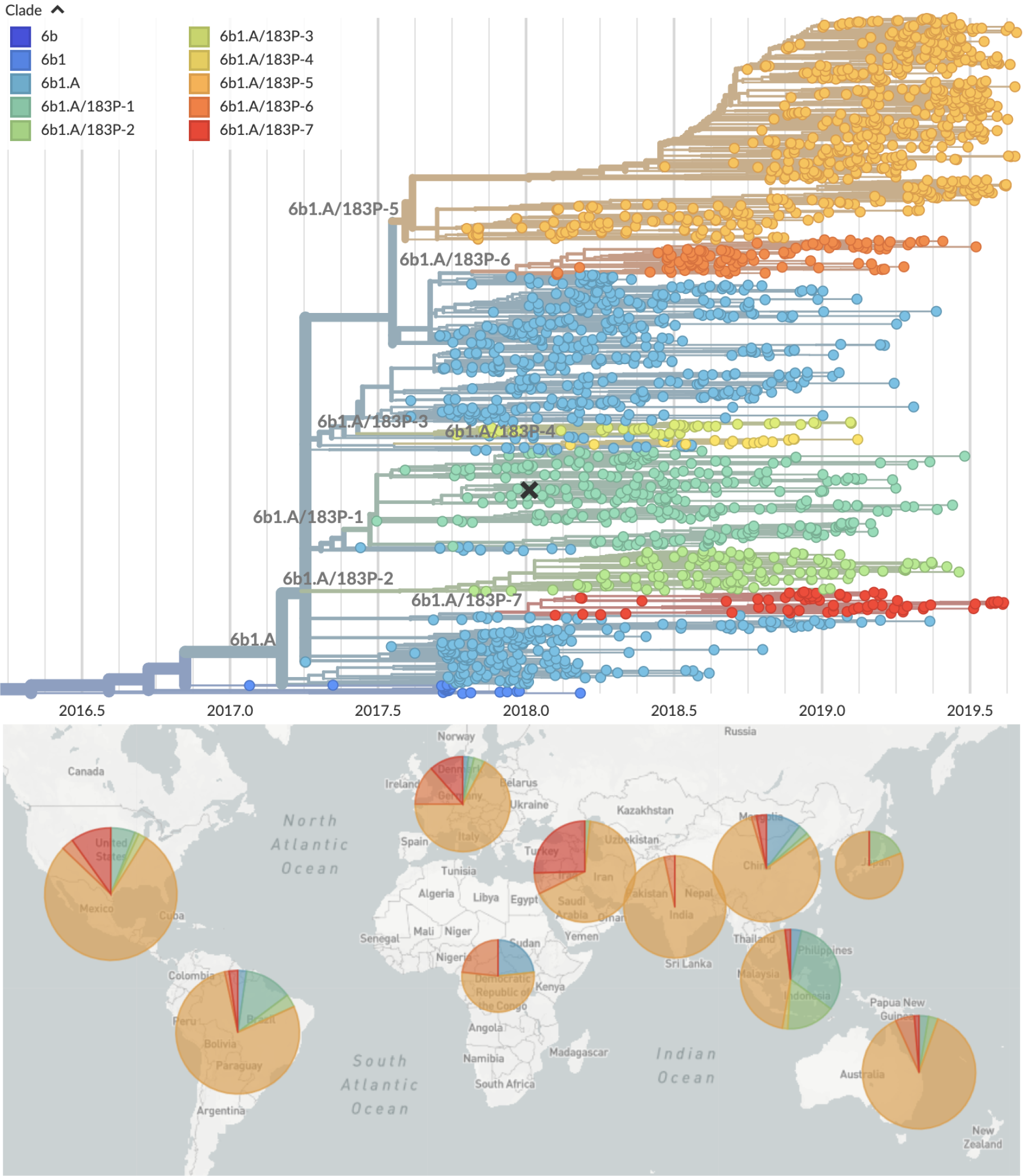
H1N1pdm phylogeny colored by clade and their geographic distribution (2019). This is a time resolved phylogeny.

**Figure 11.**
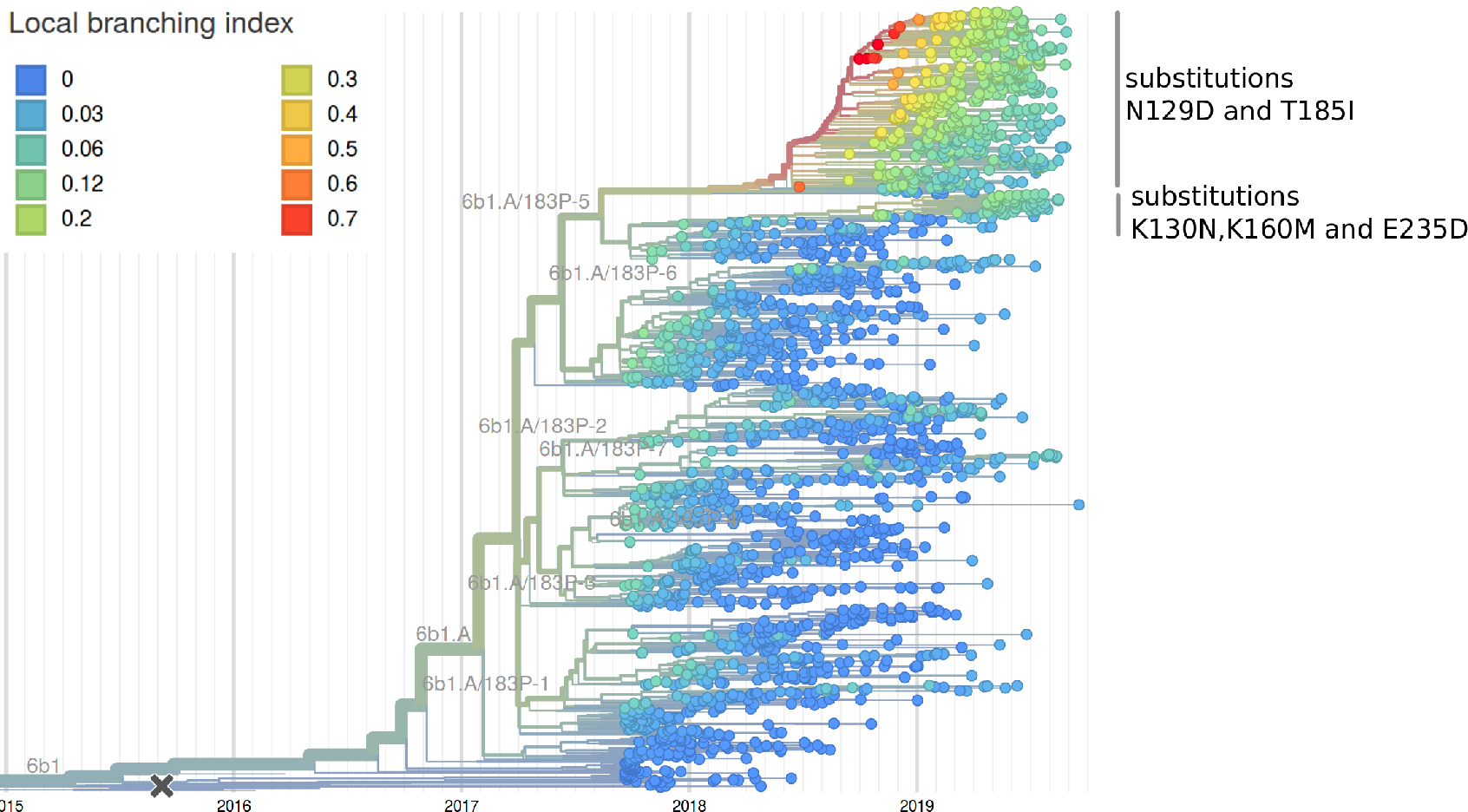
H1N1pdm phylogenies colored by LBI. The LBI accounts for both clade volume and recent expansion and has been predictive of clade success in the past. The larger subclade of P5 has the highest LBI (top), while the smaller subclade has grown rapidly in recent month. The latter might be due to an over-representation of North American sequences (see mutation HA1:130N in Fig. 13).

**Figure 12.**
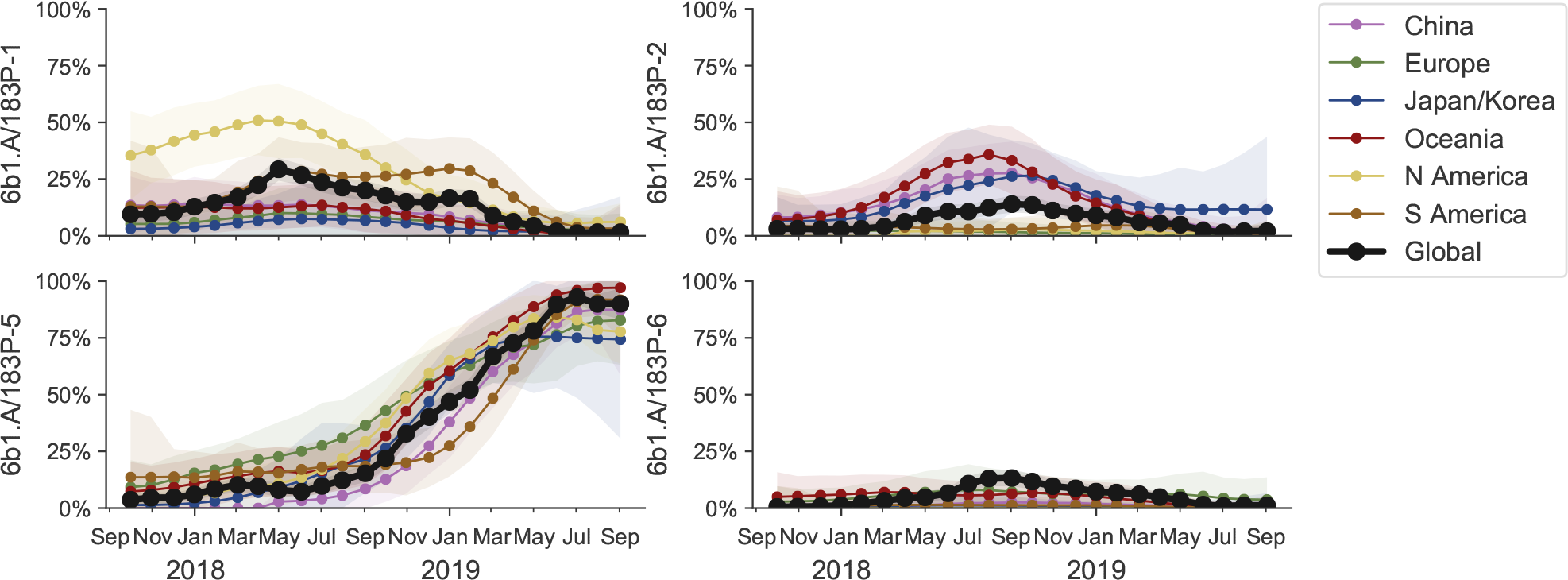
Frequency trajectories of H1N1pdm clades partitioned by clade and then by region. Clade P5 has essentially taken over, but in itself is deeply split with one subclade (substitution K130N) frequent in North America, the major subclade (T185I, N129D) prevalent elsewhere (see Figure 13). We estimate frequencies of different clades based on sample counts and collection dates of strains included in the phylogeny. We use a Brownian motion process prior to smooth frequencies from month-to-month. Transparent bands show an estimate of the 68% confidence interval based on sample counts.

This clade P5 is deeply split into a major clade with substitutions N129D and T185I and a minor clade with substitutions K160M, T216K, K130N, H296N. The relative frequencies of these two clades is best analyzed by tracking the frequencies of substitutions 129D and 130N (Fig. 13). The former is dominant in all major geographic regions, while the latter is common, but not dominant, in North America. The antigenically important substitutions N156K and N156D remain at low frequency.

**Figure 13.**
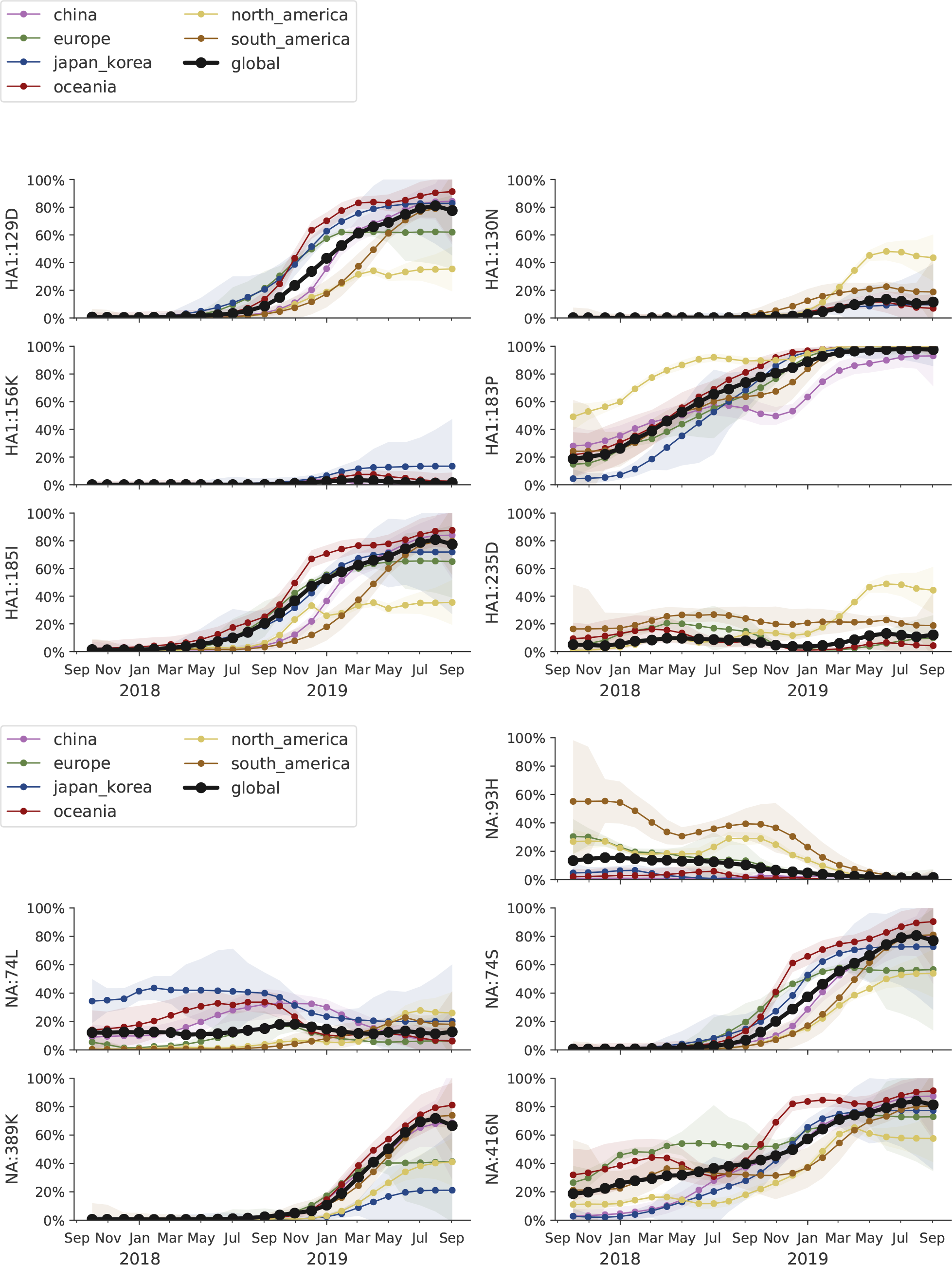
Frequency trajectories of mutations in H1N1pdm HA and NA segments partitioned by region. We estimate frequencies of different amino acid variants based on sample counts and collection dates. These estimates are based on all available data. We use a Brownian motion process prior to smooth frequencies from month-to-month. Transparent bands show an estimate of the 68% confidence interval based on sample counts.

LBI, our measure of clade success, suggests that the major subclade of P5 with substitutions N129D and T185I will continue to dominate.

Despite the rapid replacement fixation of the 183P substitution and continued diversification of H1N1pdm viruses at the HA protein level, very little antigenic evolution is apparent in HI assays. The only clear antigenic effect is associated with substitutions N156K and N156D, which result in a 16-fold titer drop. These substitutions have arisen multiple times, but are circulating at low levels and have not increased over the past 6 months. Most of the viruses with the N156K mutation fall into clade P2, most N156D variants are observed in the minor subclade of P5.

## B/Vic

Antigenically drifted deletion variants at HA1 sites 162, 163 and 164 are now dominating global circulation and have all but taken over. The double deletion variant V1A.1 had previously been circulating at high frequency in the Americas. However, over the course of 2009, the triple deletion variant V1A.3 has increased in frequency globally and is now dominating in all geographic regions. Importantly, V1A.1 and V1A.3 variants appear antigenically distinct by HI assays with 4-8 fold reductions in log2 titer in both directions.

We base our primary analysis on a set of viruses collected between Aug 2017 and Aug 2019, comprising upwards of 100 viruses per month in Nov 2018 to June 2019 (Fig. 14). Recent months are dominated by data from China and North America.

**Figure 14.**
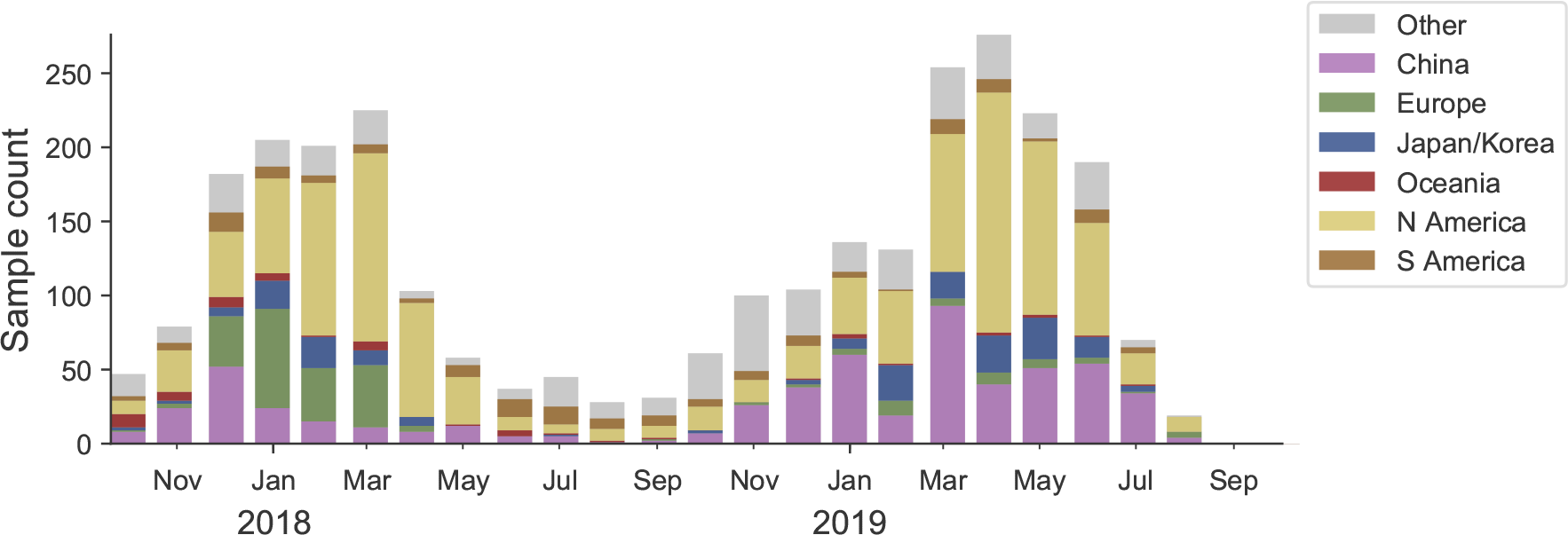
Sample counts through time and across regions. This is a stacked bar plot with the visible height of a color bar corresponding to the sample count from the respective region.

Circulating B/Vic viruses are primarily characterized be the parallel emergence of deletion variants at HA1 sites 162, 163 and 164 (Fig. 15). We label these as clades:

- Clade V1A.1: Deletions at 162 and 163. HA1 I180V. HA1 R152K.
- Clade V1A.2: Deletions at 162, 163 and 164. HA1 I180T, K209N.
- Clade V1A.3: Deletions at 162, 163 and 164. HA1 K136E.

**Figure 15.**
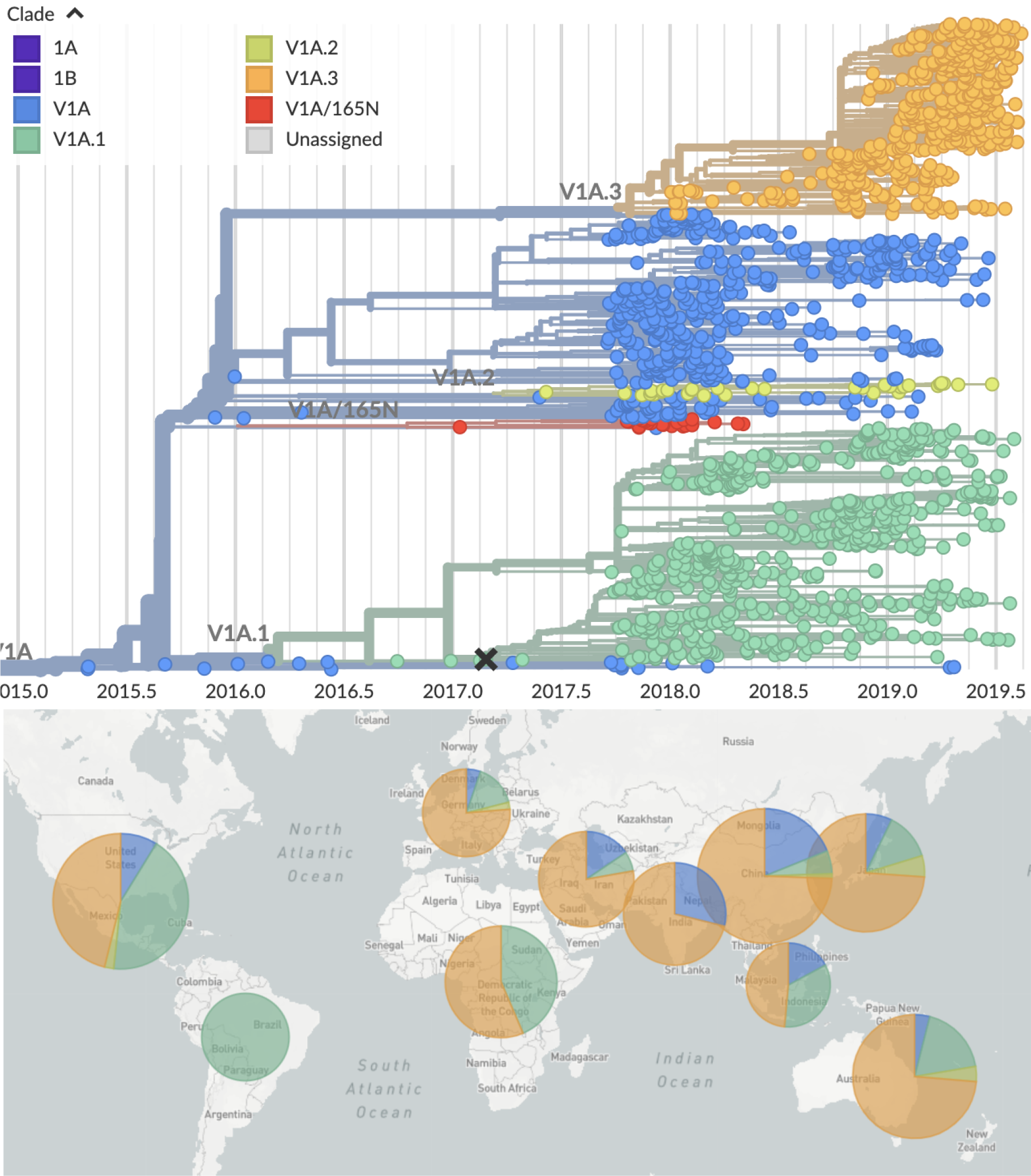
Vic phylogeny colored by clade and their geographic distribution (2019). This is a time resolved phylogeny.

Clades V1A.1 and V1A.3 have been successful with almost all 2019 viruses belonging to one or the other and V1A.3 dominating in most regions (Fig. 15).

The double deletion clade V1A.1 had risen to moderate frequency in 2018, but displayed heterogeneous frequencies across regions (Fig. 16). This clade V1A.1 is now waning in frequency. The triple deletion clade V1A.3 was initially confined to Africa, then but has rapidly in frequency in 2019, first in China and then spreading globally. Within this clade, a subclade with substitution G133R accounts for 60% of global circulation with an increasing trend (Fig. 17).

**Figure 16.**
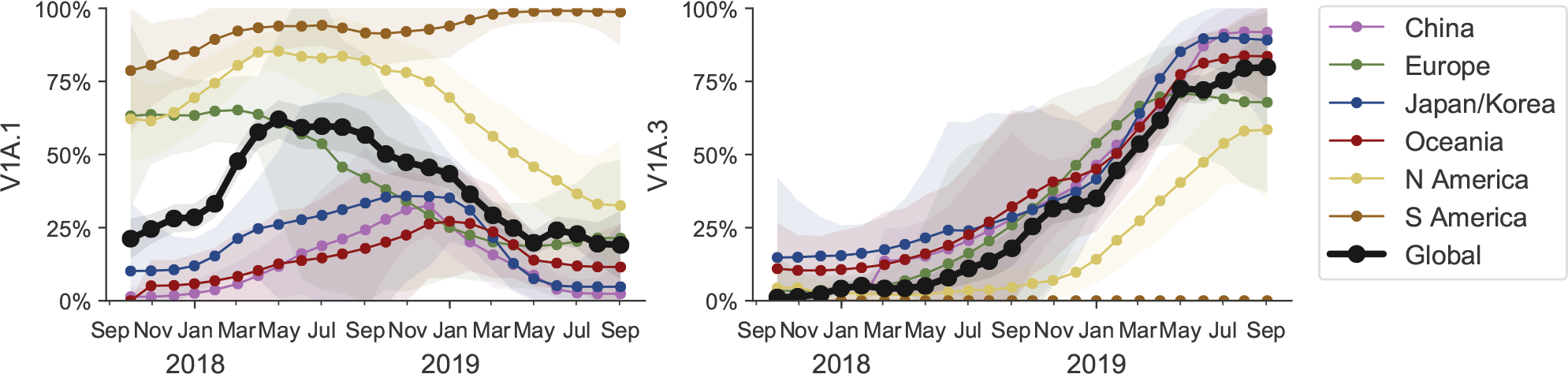
Frequency trajectories of Vic clades partitioned by clade and then by region. We estimate frequencies of different clades based on sample counts and collection dates of strains included in the phylogeny. We use a Brownian motion process prior to smooth frequencies from month-to-month. Transparent bands show an estimate of the 68% confidence interval based on sample counts.

**Figure 17.**
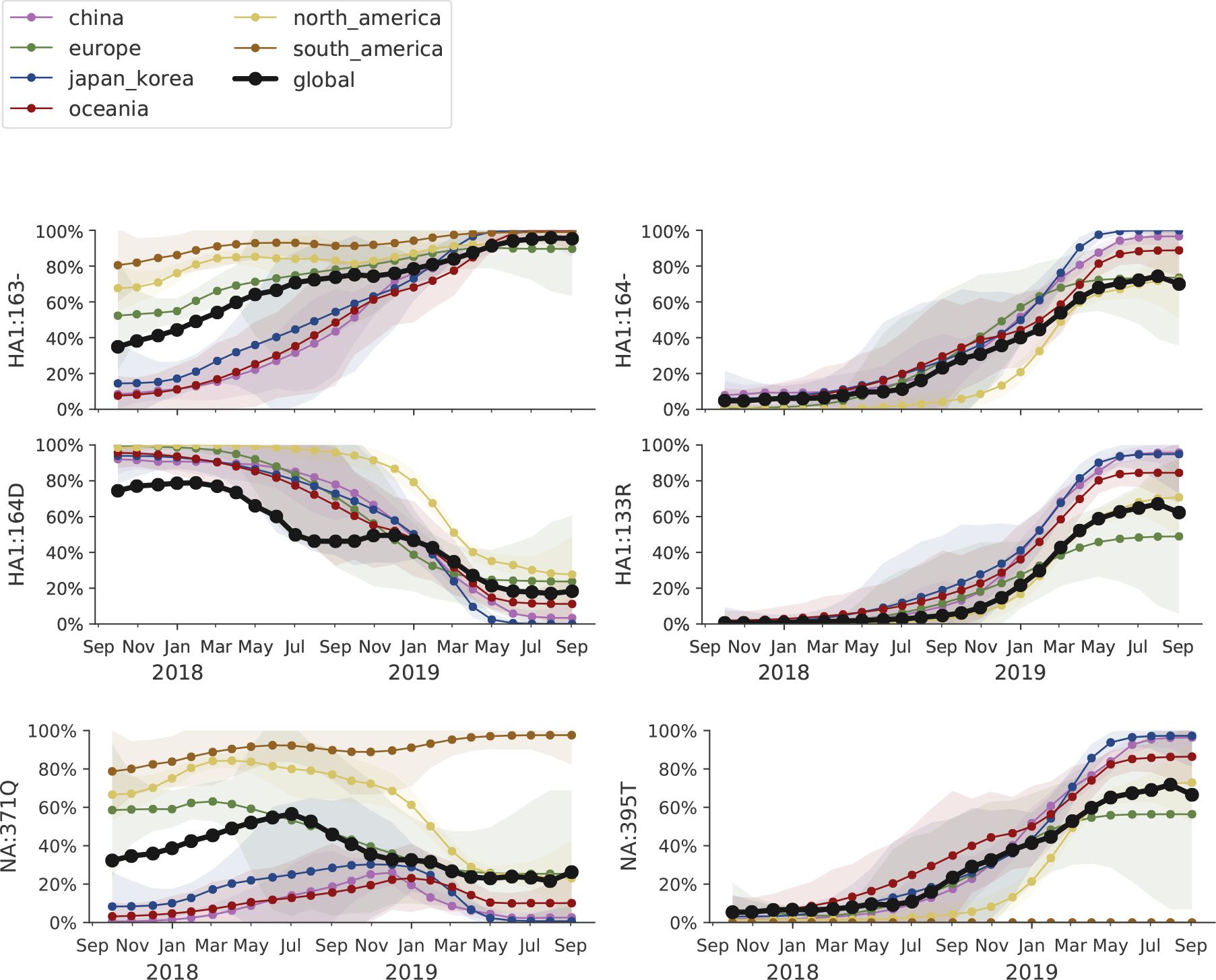
Frequency trajectories of B/Vic mutations partitioned by clade then by region. We estimate frequencies of different amino acid variants based on sample counts and collection dates. These estimates are based on all available data. We use a Brownian motion process prior to smooth frequencies from month-to-month. Transparent bands show an estimate of the 68% confidence interval based on sample counts.

HI titer data using sera raised against cell-passaged B/Colorado/6/2017 (current vaccine strain) and B/Nigeria/3352/2018 are shown on the HA phylogeny in Figure 18. Viruses from triple deletion V1A.3 clade have about 4- to 16-fold reduced titers against B/Colorado/6/2017. Analogously, viruses from double deletion clade V1A.1 are not well recognized by sera raised against B/Nigeria/3352/2018.

**Figure 18.**
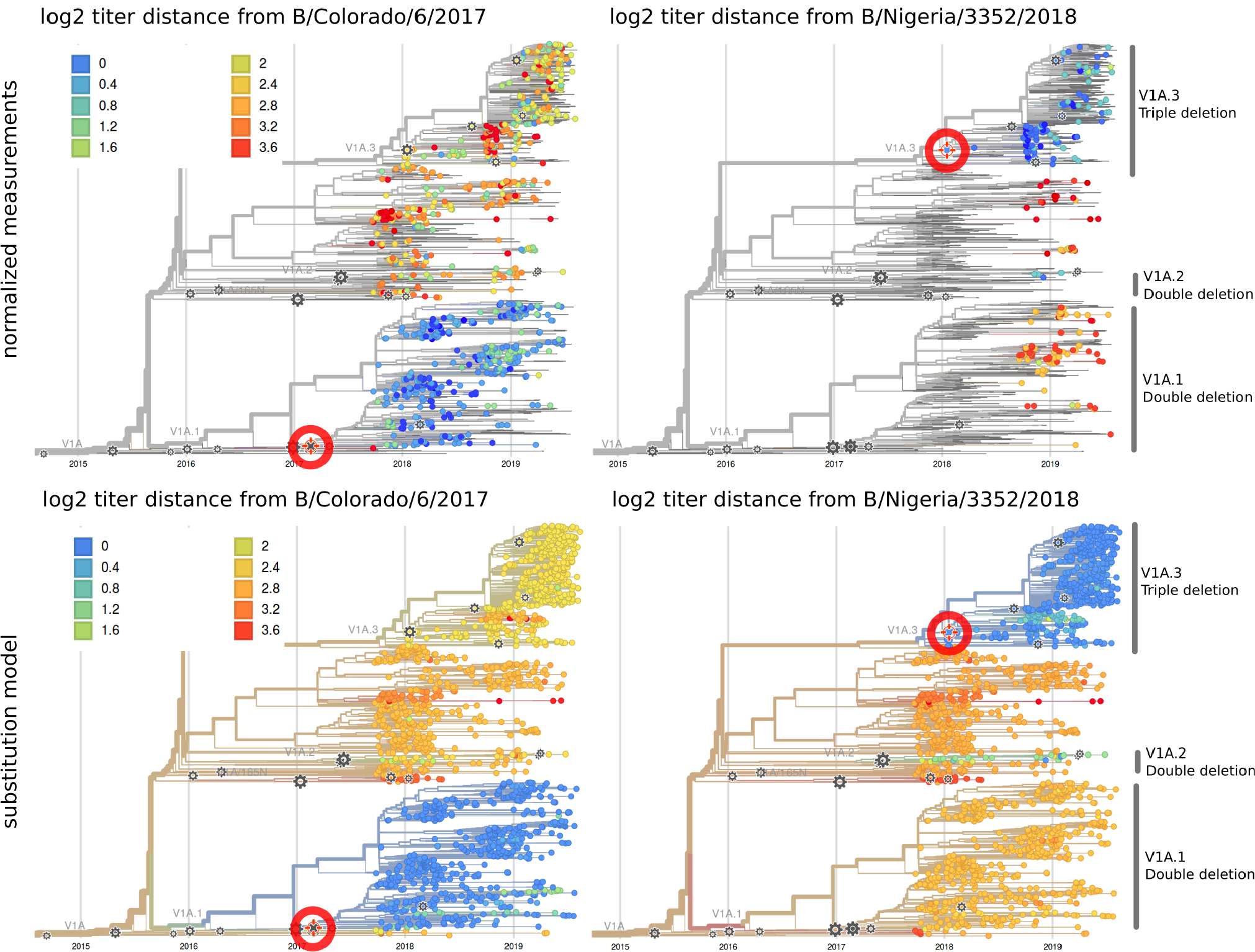
Vic phylogeny colored by antigenic distance from B/Colorado/6/2017 (left, representing V1A.1) and B/Nigeria/3352/2018 (right, representing V1A.3). While B/Colorado/6/2017 covers clade V1A.1 very well; the V1A.3 triple deletion variant (yellow clade) has about 4 to 16-fold reduced titers while isolates without a deletion have 8-fold reduced titers. B/Nigeria/3352/2018 covers both triple deletion variants well, but other viruses show 4- to 16-fold reduced titers.

## B/Yam

B/Yam has not circulated in large numbers since the Northern Hemisphere season 2017/2018 and displays relatively little amino acid variation in HA or antigenic diversity. Amino acid variants at sites 229 and 232 have begun to circulate and population is now split between 229D/232D, 229N/232D and 229D/232N variants. These variants show little sign of antigenic difference in HI assays.

We base our primary analysis on a set of viruses collected between Aug 2017 and Jul 2019, comprising between 50 and 1100 viruses per month (Fig. 19), although there are few samples after May 2018. We use all available data when estimating frequencies of mutations and weight samples appropriately by regional population size and relative sampling intensity to arrive at a putatively unbiased global frequency estimate. Phylogenetic analyses are based on a representative sample of about 2000 viruses.

**Figure 19.**
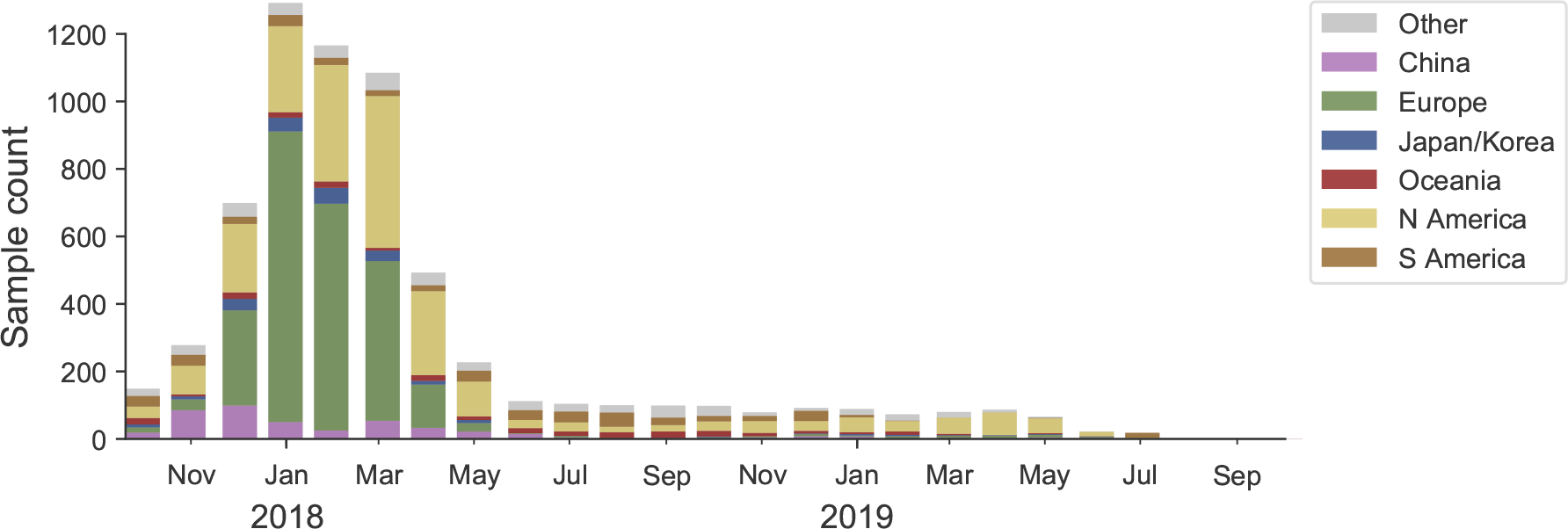
Sample counts through time and across regions. This is a stacked bar plot with the visible height of a color bar corresponding to the sample count from the respective region.

We observe very little variation among the HA segments of B/Yam viruses. Two substitutions, 229N and 232N, have started rising in frequency and are now at about 10-20% globally (Fig. 21).

**Figure 20.**
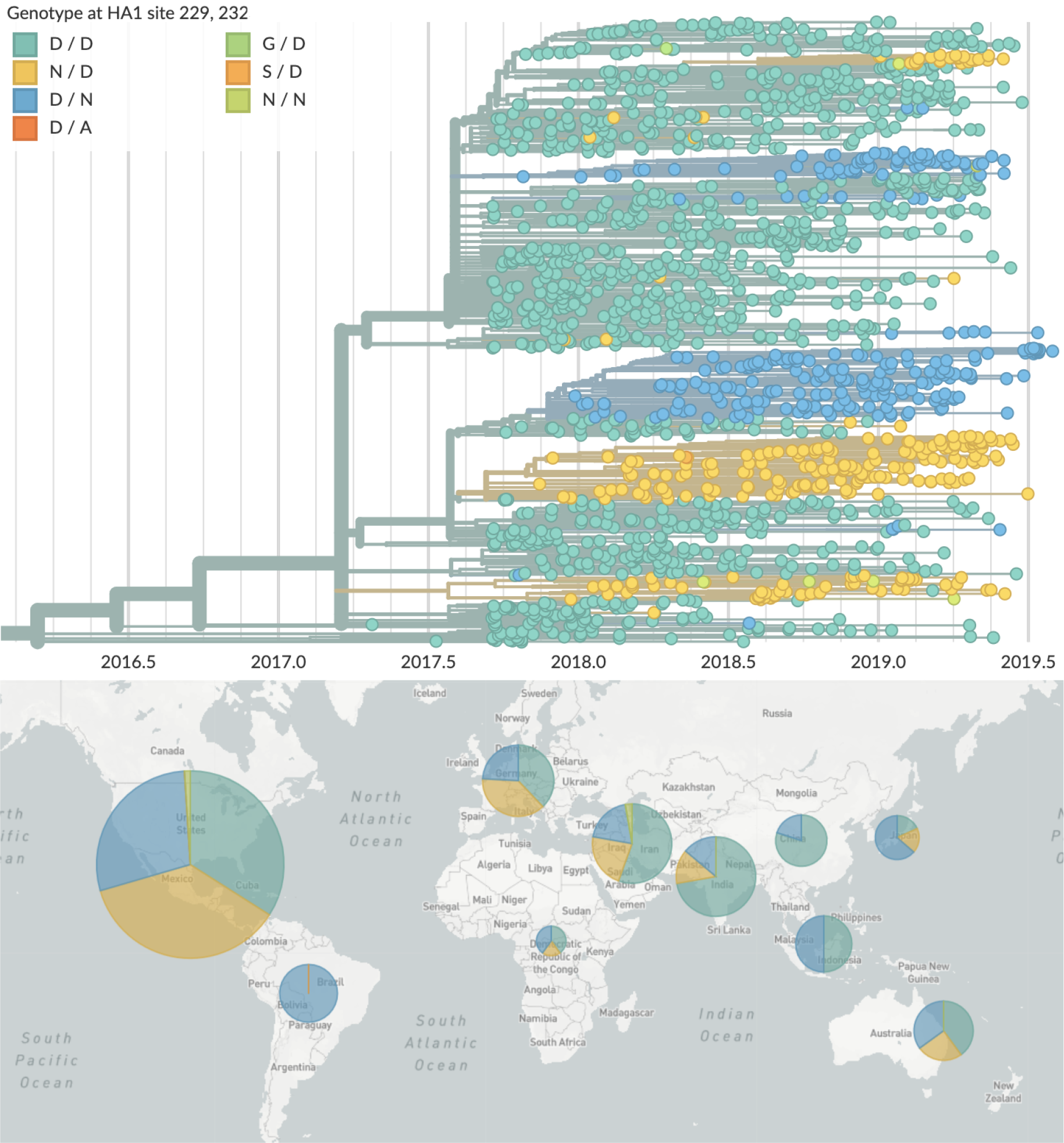
Yam phylogeny colored by mutations at sites 229 and 232 and their geographic distribution (2019). This is a time resolved phylogeny.

**Figure 21.**
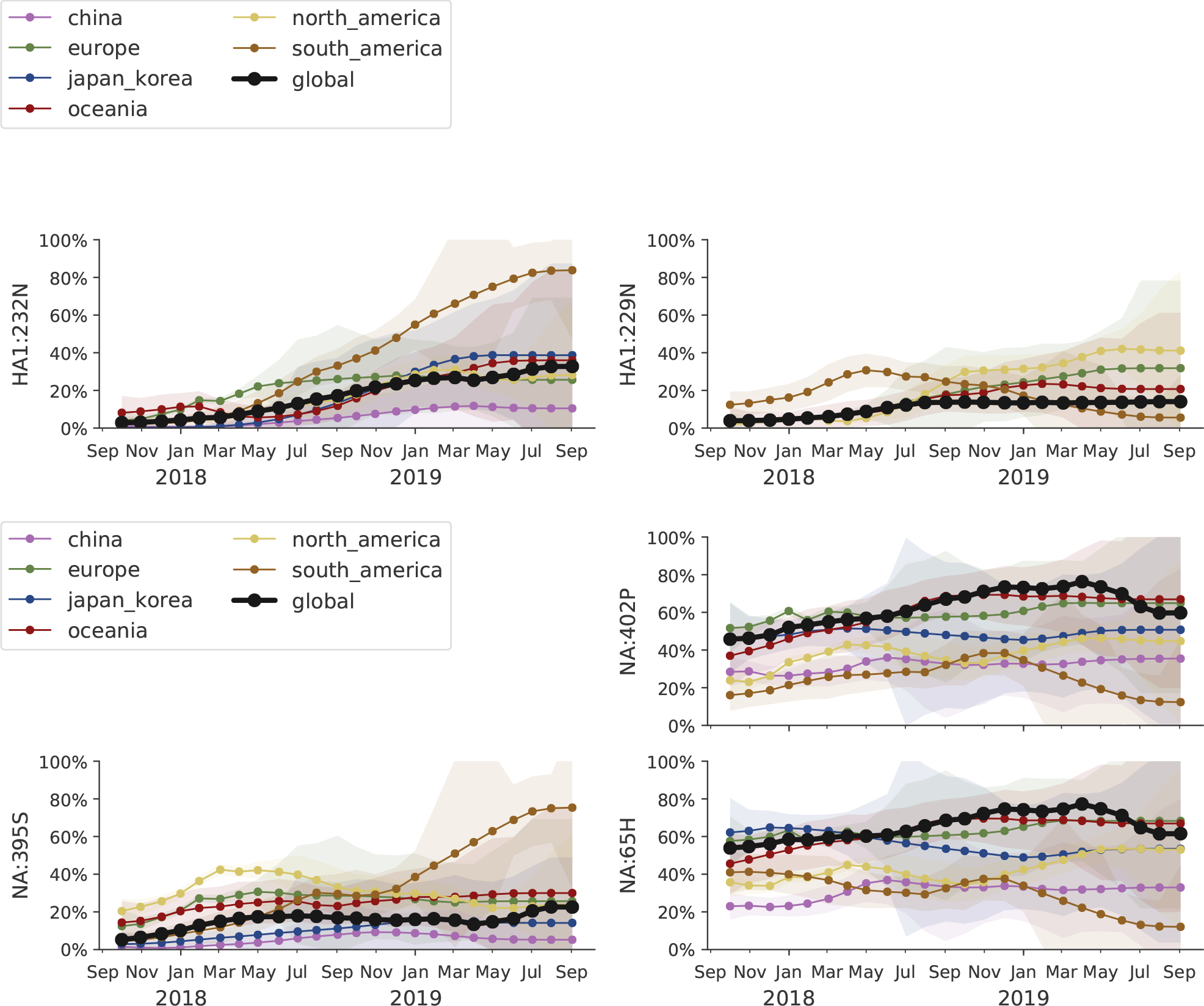
Frequency trajectories of recent mutations in B/Yam viruses. We estimate frequencies of different amino acid variants based on sample counts and collection dates. These estimates are based on all available data. We use a Brownian motion process prior to smooth frequencies from month-to-month. Transparent bands show an estimate of the 68% confidence interval based on sample counts.The almost complete lack of sequence data result in large uncertainty in these frequency estimates.

The NA segment of B/Yam had undergone a series of rapid substitutions between 2014 and 2017 but has recently become more stable (Fig. 21). The mutation 342K rose from low frequency in 2017 close to fixation. Within this clade, the nested mutations 65H and S402P are globally at about 65% frequency. Outside of this clade, A395S is common in North America and Europe (Fig. 21). Site 342 has changed twice in recent years: D342N followed by N342K.

## Acknowledgments

We thank the Influenza Division at the US Centers for Disease Control and Prevention, the Victorian Infectious Diseases Reference Laboratory at the Australian Peter Doherty Institute for Infection and Immunity, the Influenza Virus Research Center at the Japan National Institute of Infectious Diseases, the Crick Worldwide Influenza Centre at the UK Francis Crick Institute for data sharing and feedback. We thank David Wentworth, Rebecca Kondor and Vivien Dugan for insight regarding analysis directions.

We gratefully acknowledge the authors, originating and submitting laboratories of the sequences from the GISAID EpiFlu Database [3] on which this research is based. The list is detailed below:

WHO Collaborating Centre for Reference and Research on Influenza, Victorian Infectious Diseases Reference Laboratory, Australia; WHO Collaborating Centre for Reference and Research on Influenza, Chinese National Influenza Center, China; WHO Collaborating Centre for Reference and Research on Influenza, National Institute of Infectious Diseases, Japan; The Crick Worldwide Influenza Centre, The Francis Crick Institute, United Kingdom; WHO Collaborating Centre for the Surveillance, Epidemiology and Control of Influenza, Centers for Disease Control and Prevention, United States; ADImmune Corporation, Taiwan; ADPH Bureau of Clinical Laboratories, United States; Aichi Prefectural Institute of Public Health, Japan; Akershus University Hospital, Norway; Akita Research Center for Public Health and Environment, Japan; Alabama State Laboratory, United States; Alaska State Public Health Laboratory, United States; Alaska State Virology Lab, United States; Aomori Prefectural Institute of Public Health and Environment, Japan; Aristotelian University of Thessaloniki, Greece; Arizona Department of Health Services, United States; Arkansas Children’s Hospital, United States; Arkansas Department of Health, United States; Auckland Healthcare, New Zealand; Auckland Hospital, New Zealand; Austin Health, Australia; Baylor College of Medicine, United States; California Department of Health Services, United States; Canberra Hospital, Australia; Cantacuzino Institute, Romania; Canterbury Health Services, New Zealand; Caribbean Epidemiology Center, Trinidad and Tobago; CDC GAP Nigeria, Nigeria; CDC-Kenya, Kenya; CEMIC University Hospital, Argentina; CENETROP, Bolivia, Plurinationial State of; Center for Disease Control, Taiwan; Center for Public Health and Environment, Hiroshima Prefectural Technology Research Institute, Japan; Central Health Laboratory, Mauritius; Central Laboratory of Public Health, Paraguay; Central Public Health Laboratory, Ministry of Health, Oman; Central Public Health Laboratory, Palestinian Territory; Central Public Health Laboratory, Papua New Guinea; Central Research Institute for Epidemiology, Russian Federation; Centre for Diseases Control and Prevention, Armenia; Centre for Infections, Health Protection Agency, United Kingdom; Centre Pasteur du Cameroun, Cameroon; Chiba City Institute of Health and Environment, Japan; Chiba Prefectural Institute of Public Health, Japan; Childrens Hospital Westmead, Australia; Chuuk State Hospital, Micronesia, Federated States of; City of El Paso Dept of Public Health, United States; Clinical Virology Unit, CDIM, Australia; Colorado Department of Health Lab, United States; Connecticut Department. of Public Health, United States; Contiguo a Hospital Rosales, El Salvador; Croatian Institute of Public Health, Croatia; CRR virus Influenza region Sud, France; CRR virus Influenza region Sud, Guyana; CSL Ltd, United States; Dallas County Health and Human Services, United States; DC Public Health Lab, United States; Delaware Public Health Lab, United States; Departamento de Laboratorio de Salud Publica, Uruguay; Department of Virology, Medical University Vienna, Austria; Disease Investigation Centre Wates (BBVW), Australia; Drammen Hospital / Vestreviken HF, Norway; Ehime Prefecture Institute of Public Health and Environmental Science, Japan; Erasmus Medical Center, Netherlands; Erasmus University of Rotterdam, Netherlands; Ethiopian Health and Nutrition Research Institute (EHNRI), Ethiopia; Evanston Hospital and NorthShore University, United States; Facultad de Medicina, Spain; Fiji Centre for Communicable Disease Control, Fiji; Florida Department of Health, United States; Fukui Prefectural Institute of Public Health, Japan; Fukuoka City Institute for Hygiene and the Environment, Japan; Fukuoka Institute of Public Health and Environmental Sciences, Japan; Fukushima Prefectural Institute of Public Health, Japan; Gart Naval General Hospital, United Kingdom; Georgia Public Health Laboratory, United States; Gifu Municipal Institute of Public Health, Japan; Gifu Prefectural Institute of Health and Environmental Sciences, Japan; Government Virus Unit, Hong Kong; Gunma Prefectural Institute of Public Health and Environmental Sciences, Japan; Hamamatsu City Health Environment Research Center, Japan; Haukeland University Hospital, Dept. of Microbiology, Norway; Headquarters British Gurkhas Nepal, Nepal; Health Forde, Department of Microbiology, Norway; Health Protection Agency, United Kingdom; Health Protection Inspectorate, Estonia; Hellenic Pasteur Institute, Greece; Hiroshima City Institute of Public Health, Japan; Hokkaido Institute of Public Health, Japan; Hopital Cantonal Universitaire de Geneves, Switzerland; Hopital Charles Nicolle, Tunisia; Hospital Clinic de Barcelona, Spain; Hospital Universitari Vall d’Hebron, Spain; Houston Department of Health and Human Services, United States; Hyogo Prefectural Institute of Public Health and Consumer Sciences, Japan; Ibaraki Prefectural Institute of Public Health, Japan; Illinois Department of Public Health, United States; Indiana State Department of Health Laboratories, United States; Infectology Center of Latvia, Latvia; Innlandet Hospital Trust, Division Lillehammer, Department for Microbiology, Norway; INSA National Institute of Health Portugal, Portugal; Institut National d’Hygiene, Morocco; Institut Pasteur d’Algerie, Algeria; Institut Pasteur de Dakar, Senegal; Institut Pasteur de Madagascar, Madagascar; Institut Pasteur in Cambodia, Cambodia; Institut Pasteur New Caledonia, New Caledonia; Institut Pasteur, France; Institut Pasteur, Saudi Arabia; Institut Penyelidikan Perubatan, Malaysia; Institute National D’Hygiene, Togo; Institute of Environmental Science and Research, New Zealand; Institute of Environmental Science and Research, Tonga; Institute of Epidemiology and Infectious Diseases, Ukraine; Institute of Epidemiology Disease Control and Research, Bangladesh; Institute of Immunology and Virology Torlak, Serbia; Institute of Medical and Veterinary Science (IMVS), Australia; Institute of Public Health, Serbia; Institute of Public Health, Albania; Institute of Public Health, Montenegro; Institute Pasteur du Cambodia, Cambodia; Instituto Adolfo Lutz, Brazil; Instituto Conmemorativo Gorgas de Estudios de la Salud, Panama; Instituto de Salud Carlos III, Spain; Instituto de Salud Publica de Chile, Chile; Instituto Nacional de Enfermedades Infecciosas, Argentina; Instituto Nacional de Higiene Rafael Rangel, Venezuela, Bolivia; Instituto Nacional de Laboratoriosde Salud (INLASA), Bolivia; Instituto Nacional de Salud de Columbia, Colombia; Instituto Nacional de Saude, Portugal; Iowa State Hygienic Laboratory, United States; IRSS, Burkina Faso; Ishikawa Prefectural Institute of Public Health and Environmental Science, Japan; ISS, Italy; Istanbul University, Turkey; Istituto Superiore di Sanit, Italy; Ivanovsky Research Institute of Virology RAMS, Russian Federation; Jiangsu Provincial Center for Disease Control and Prevention, China; John Hunter Hospital, Australia; Kagawa Prefectural Research Institute for Environmental Sciences and Public Health, Japan; Kagoshima Prefectural Institute for Environmental Research and Public Health, Japan; Kanagawa Prefectural Institute of Public Health, Japan; Kansas Department of Health and Environment, United States; Kawasaki City Institute of Public Health, Japan; Kentucky Division of Laboratory Services, United States; Kitakyusyu City Institute of Enviromental Sciences, Japan; Kobe Institute of Health, Japan; Kochi Public Health and Sanitation Institute, Japan; Kumamoto City Environmental Research Center, Japan; Kumamoto Prefectural Institute of Public Health and Environmental Science, Japan; Kyoto City Institute of Health and Environmental Sciences, Japan; Kyoto Prefectural Institute of Public Health and Environment, Japan; Laboratoire National de Sante Publique, Haiti; Laboratoire National de Sante, Luxembourg; Laboratrio Central do Estado do Paran, Brazil; Laboratorio Central do Estado do Rio de Janeiro, Brazil; Laboratorio de Investigacion / Centro de Educacion Medica y Amistad Dominico Japones (CEMADOJA), Dominican Republic; Laboratorio De Saude Publico, Macao; Laboratorio de Virologia, Direccion de Microbiologia, Nicaragua; Laboratorio de Virus Respiratorio, Mexico; Laboratorio Nacional de Influenza, Costa Rica; Laboratorio Nacional De Salud Guatemala, Guatemala; Laboratorio Nacional de Virologia, Honduras; Laboratory Directorate, Jordan; Laboratory for Virology, National Institute of Public Health, Slovenia; Laboratory of Influenza and ILI, Belarus; LACEN/RS - Laboratrio Central de Sade Pblica do Rio Grande do Sul, Brazil; Landspitali - University Hospital, Iceland; Lithuanian AIDS Center Laboratory, Lithuania; Los Angeles Quarantine Station, CDC Quarantine Epidemiology and Surveillance Team, United States; Louisiana Department of Health and Hospitals, United States; Maine Health and Environmental Testing Laboratory, United States; Malbran, Argentina; Marshfield Clinic Research Foundation, United States; Maryland Department of Health and Mental Hygiene, United States; Massachusetts Department of Public Health, United States; Mater Dei Hospital, Malta; Medical Research Institute, Sri Lanka; Medical University Vienna, Austria; Melbourne Pathology, Australia; Michigan Department of Community Health, United States; Mie Prefecture Health and Environment Research Institute, Japan; Mikrobiologisk laboratorium, Sykehuset i Vestfold, Norway; Ministry of Health and Population, Egypt; Ministry of Health of Ukraine, Ukraine; Ministry of Health, Bahrain; Ministry of Health, Kiribati; Ministry of Health, Lao, People’s Democratic Republic; Ministry of Health, NIHRD, Indonesia; Ministry of Health, Oman; Minnesota Department of Health, United States; Mississippi Public Health Laboratory, United States; Missouri Department. of Health and Senior Services, United States; Miyagi Prefectural Institute of Public Health and Environment, Japan; Miyazaki Prefectural Institute for Public Health and Environment, Japan; Molde Hospital, Laboratory for Medical Microbiology, Norway; Molecular Diagnostics Unit, United Kingdom; Monash Medical Centre, Australia; Montana Laboratory Services Bureau, United States; Montana Public Health Laboratory, United States; Nagano City Health Center, Japan; Nagano Environmental Conservation Research Institute, Japan; Nagoya City Public Health Research Institute, Japan; Nara Prefectural Institute for Hygiene and Environment, Japan; National Center for Communicable Diseases, Mongolia; National Center for Laboratory and Epidemiology, Laos; National Centre for Disease Control (NCDC), Mongolia; National Centre for Disease Control and Public Health, Georgia; National Centre for Preventive Medicine, Moldova, Republic of; National Centre for Scientific Services for Virology and Vector Borne Diseases, Fiji; National Health Laboratory, Japan; National Health Laboratory, Myanmar; National Influenza Center French Guiana and French Indies, French Guiana; National Influenza Center, Brazil; National Influenza Center, Mongolia; National Influenza Centre for Northern Greece, Greece; National Influenza Centre of Iraq, Iraq; National Influenza Lab, Tanzania, United Republic of; National Influenza Reference Laboratory, Nigeria; National Insitut of Hygien, Morocco; National Institute for Biological Standards and Control (NIBSC), United States; National Institute for Communicable Disease, South Africa; National Institute for Health and Welfare, Finland; National Institute of Health Research and Development, Indonesia; National Institute of Health, Korea, Republic of; National Institute of Health, Pakistan; National Institute of Hygien, Morocco; National Institute of Hygiene and Epidemiology, Vietnam; National Institute of Public Health - National Institute of Hygiene, Poland; National Institute of Public Health, Czech Republic; National Institute of Virology, India; National Microbiology Laboratory, Health Canada, Canada; National Public Health Institute of Slovakia, Slovakia; National Public Health Laboratory, Cambodia; National Public Health Laboratory, Ministry of Health, Singapore, Singapore; National Public Health Laboratory, Nepal; National Public Health Laboratory, Singapore; National Reference Laboratory, Kazakhstan; National University Hospital, Singapore; National Virology Laboratory, Center Microbiological Investigations, Kyrgyzstan; National Virus Reference Laboratory, Ireland; Naval Health Research Center, United States; Nebraska Public Health Lab, United States; Nevada State Health Laboratory, United States; New Hampshire Public Health Laboratories, United States; New Jersey Department of Health and Senior Services, United States; New Mexico Department of Health, United States; New York City Department of Health, United States; New York Medical College, United States; New York State Department of Health, United States; Nicosia General Hospital, Cyprus; Niigata City Institute of Public Health and Environment, Japan; Niigata Prefectural Institute of Public Health and Environmental Sciences, Japan; Niigata University, Japan; Nordlandssykehuset, Norway; North Carolina State Laboratory of Public Health, United States; North Dakota Department of Health, United States; Norwegian Institute of Public Health, Norway; Norwegian Institute of Public Health, Svalbard and Jan Mayen; Ohio Department of Health Laboratories, United States; Oita Prefectural Institute of Health and Environment, Japan; Okayama Prefectural Institute for Environmental Science and Public Health, Japan; Okinawa Prefectural Institute of Health and Environment, Japan; Oklahoma State Department of Health, United States; Ontario Agency for Health Protection and Promotion (OAHPP), Canada; Oregon Public Health Laboratory, United States; Osaka City Institute of Public Health and Environmental Sciences, Japan; Osaka Prefectural Institute of Public Health, Japan; Oslo University Hospital, Ulleval Hospital, Dept. of Microbiology, Norway; Ostfold Hospital - Fredrikstad, Dept. of Microbiology, Norway; Oswaldo Cruz Institute - FIOCRUZ - Laboratory of Respiratory Viruses and Measles (LVRS), Brazil; Papua New Guinea Institute of Medical Research, Papua New Guinea; Pasteur Institut of Cote d’Ivoire, Cote d’Ivoire; Pasteur Institute, Influenza Laboratory, Vietnam; Pathwest QE II Medical Centre, Australia; Pennsylvania Department of Health, United States; Prince of Wales Hospital, Australia; Princess Margaret Hospital for Children, Australia; Public Health Laboratory Services Branch, Centre for Health Protection, Hong Kong; Public Health Laboratory, Barbados; Puerto Rico Department of Health, Puerto Rico; Qasya Diagnostic Services Sdn Bhd, Brunei; Queensland Health Scientific Services, Australia; Refik Saydam National Public Health Agency, Turkey; Regent Seven Seas Cruises, United States; Royal Victoria Hospital, United Kingdom; Republic Institute for Health Protection, Macedonia, the former Yogoslav Republic of; Republic of Nauru Hospital, Nauru; Research Institute for Environmental Sciences and Public Health of Iwate Prefecture, Japan; Research Institute of Tropical Medicine, Philippines; Rhode Island Department of Health, United States; RIVM National Institute for Public Health and Environment, Netherlands; Robert-Koch-Institute, Germany; Royal Chidrens Hospital, Australia; Royal Darwin Hospital, Australia; Royal Hobart Hospital, Australia; Royal Melbourne Hospital, Australia; Russian Academy of Medical Sciences, Russian Federation; Rwanda Biomedical Center, National Reference Laboratory, Rwanda; Saga Prefectural Institute of Public Health and Pharmaceutical Research, Japan; Sagamihara City Laboratory of Public Health, Japan; Saitama City Institute of Health Science and Research, Japan; Saitama Institute of Public Health, Japan; Sakai City Institute of Public Health, Japan; San Antonio Metropolitan Health, United States; Sandringham, National Institute for Communicable D, South Africa; Sapporo City Institute of Public Health, Japan; Scientific Institute of Public Health, Belgium; Seattle and King County Public Health Lab, United States; Sendai City Institute of Public Health, Japan; Servicio de Microbiologa Clnica Universidad de Navarra, Spain; Servicio de Microbiologa Complejo Hospitalario de Navarra, Spain; Servicio de Microbiologa Hospital Central Universitario de Asturias, Spain; Servicio de Microbiologa Hospital Donostia, Spain; Servicio de Microbiologa Hospital Meixoeiro, Spain; Servicio de Microbiologa Hospital Miguel Servet, Spain; Servicio de Microbiologa Hospital Ramn y Cajal, Spain; Servicio de Microbiologa Hospital San Pedro de Alcntara, Spain; Servicio de Microbiologa Hospital Santa Mara Nai, Spain; Servicio de Microbiologa Hospital Universitario de Gran Canaria Doctor Negrn, Spain; Servicio de Microbiologa Hospital Universitario Son Espases, Spain; Servicio de Microbiologa Hospital Virgen de la Arrixaca, Spain; Servicio de Microbiologa Hospital Virgen de las Nieves, Spain; Servicio de Virosis Respiratorias INEI-ANLIS Carlos G. Malbran, Argentina; Shiga Prefectural Institute of Public Health, Japan; Shimane Prefectural Institute of Public Health and Environmental Science, Japan; Shizuoka City Institute of Environmental Sciences and Public Health, Japan; Shizuoka Institute of Environment and Hygiene, Japan; Singapore General Hospital, Singapore; Sorlandet Sykehus HF, Dept. of Medical Microbiology, Norway; South Carolina Department of Health, United States; South Dakota Public Health Lab, United States; Southern Nevada Public Health Lab, United States; Spokane Regional Health District, United States; St. Judes Childrens Research Hospital, United States; St. Olavs Hospital HF, Dept. of Medical Microbiology, Norway; State Agency, Infectology Center of Latvia, Latvia; State of Hawaii Department of Health, United States; State of Idaho Bureau of Laboratories, United States; State Research Center of Virology and Biotechnology Vector, Russian Federation; Statens Serum Institute, Denmark; Stavanger Universitetssykehus, Avd. for Medisinsk Mikrobiologi, Norway; Subdireccion General de Epidemiologia y Vigilancia de la Salud, Spain; Subdireccin General de Epidemiologa y Vigilancia de la Salud, Spain; Swedish Institute for Infectious Disease Control, Sweden; Swedish National Institute for Communicable Disease Control, Sweden; Taiwan CDC, Taiwan; Tan Tock Seng Hospital, Singapore; Tehran University of Medical Sciences, Iran; Tennessee Department of Health Laboratory-Nashville, United States; Texas Childrens Hospital, United States; Texas Department of State Health Services, United States; Thai National Influenza Center, Thailand; Thailand MOPH-U.S. CDC Collaboration (IEIP), Thailand; The Nebraska Medical Center, United States; Tochigi Prefectural Institute of Public Health and Environmental Science, Japan; Tokushima Prefectural Centre for Public Health and Environmental Sciences, Japan; Tokyo Metropolitan Institute of Public Health, Japan; Tottori Prefectural Institute of Public Health and Environmental Science, Japan; Toyama Institute of Health, Japan; U.S. Air Force School of Aerospace Medicine, United States; U.S. Naval Medical Research Unit No.3, Egypt; Uganda Virus Research Institute (UVRI), National Influenza Center, Uganda; Universidad de Valladolid, Spain; Universit Cattolica del Sacro Cuore, Italy; Universitetssykehuset Nord-Norge HF, Norway; University Malaya, Malaysia; University of Florence, Italy; University of Genoa, Italy; University of Ghana, Ghana; University of Michigan SPH EPID, United States; University of Parma, Italy; University of Perugia, Italy; University of Pittsburgh Medical Center Microbiology Lab, United States; University of Sarajevo, Bosnia and Herzegovina; University of Sassari, Italy; University of the West Indies, Jamaica; University of Vienna, Austria; University of Virginia, Medical Labs/Microbiology, United States; University Teaching Hospital, Zambia; UPMC-CLB Dept of Microbiology, United States; US Army Medical Research Unit - Kenya (USAMRU-K), GEIS Human Influenza Program, Kenya; USAMC-AFRIMS Department of Virology, Cambodia; Utah Department of Health, United States; Utah Public Health Laboratory, United States; Utsunomiya City Institute of Public Health and Environment Science, Japan; VACSERA, Egypt; Vermont Department of Health Laboratory, United States; Victorian Infectious Diseases Reference Laboratory, Australia; Virginia Division of Consolidated Laboratories, United States; Wakayama City Institute of Public Health, Japan; Wakayama Prefectural Research Center of Environment and Public Health, Japan; Washington State Public Health Laboratory, United States; West Virginia Office of Laboratory Services, United States; Westchester County Department of Laboratories and Research, United States; Westmead Hospital, Australia; WHO National Influenza Centre Russian Federation, Russian Federation; WHO National Influenza Centre, National Institute of Medical Research (NIMR), Thailand; WHO National Influenza Centre, Norway; Wisconsin State Laboratory of Hygiene, United States; Wyoming Public Health Laboratory, United States; Yamagata Prefectural Institute of Public Health, Japan; Yamaguchi Prefectural Institute of Public Health and Environment, Japan; Yamanashi Institute for Public Health, Japan; Yap State Hospital, Micronesia; Yokohama City Institute of Health, Japan; Yokosuka Institute of Public Health, Japan

## Host age distributions

**Figure 22.**
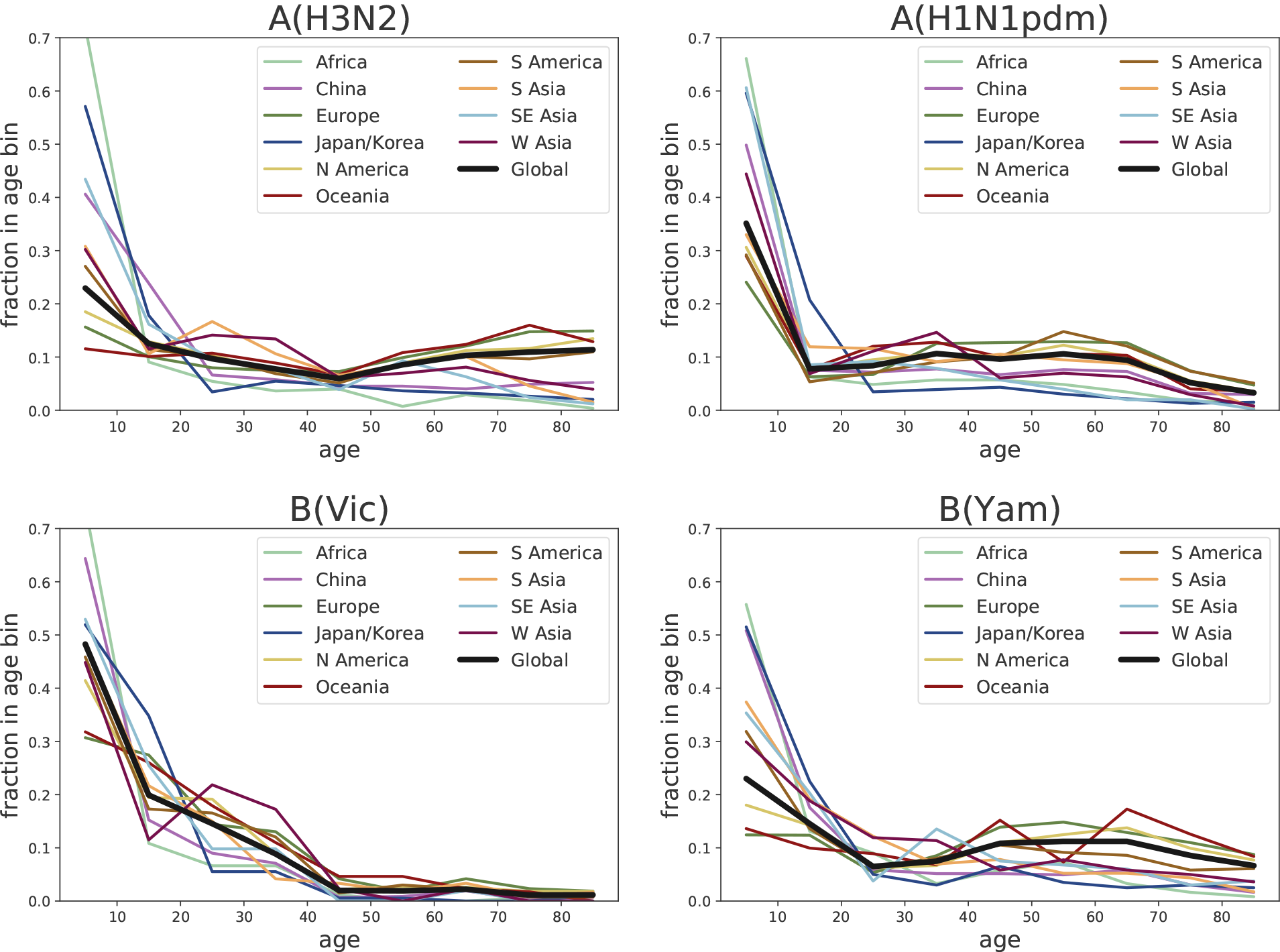
Each panel show the fraction of sequences sampled from hosts in a 10 year age bin for different geographic regions. These distributions are confounded by reporting bias, but the differences between different lineages are big enough that the overall picture is unlikely changed by these biases. As expected, H1N1pdm sequences are more frequently sampled from young children compared to H3N2. The biggest difference is observed between the two influenza B lineages: B/Vic viruses were mostly sampled from young individuals, in sharp contrast to B/Yam which is frequently isolated from patients above 40y of age, in particular in Europe, North America, and Oceania.

## Notes

#### Summary of Updates

The previous version omitted acknowledgments of sequence data providers. This version has "Acknowledgements" updated with an explicit list of laboratories that shared sequence data via the GISAID database.

https://nextstrain.org/flu

## References

1. Neher RA, Bedford T (2015) nextflu: real-time tracking of seasonal influenza virus evolution in humans. Bioinformatics 31: 3546–3548.

2. Hadfield J, Megill C, Bell SM, Huddleston J, Potter B, et al. (2018) Nextstrain: real-time tracking of pathogen evolution. Bioinformatics 34: 4121–4123.

3. Shu Y, McCauley J (2017) Gisaid: Global initiative on sharing all influenza data – from vision to reality. Euro-surveillance 22.

4. Neher RA, Bedford T, Daniels RS, Russell CA, Shraiman BI (2016) Prediction, dynamics, and visualization of antigenic phenotypes of seasonal influenza viruses. Proc Natl Acad Sci USA 113: E1701–E1709.

5. Neher RA, Russell CA, Shraiman BI (2014) Predicting evolution from the shape of genealogical trees. eLife 3: e03568.

